# A proteogenomic pipeline for the analysis of protein biosynthesis errors in the human pathogen *Candida albicans*

**DOI:** 10.1101/2023.10.31.564356

**Authors:** Inês Correia, Carla Oliveira, Andreia Reis, Ana Rita Guimarães, Susana Aveiro, Pedro Domingues, Ana Rita Bezerra, Rui Vitorino, Gabriela Moura, Manuel A. S. Santos

**Affiliations:** Institute of Biomedicine (iBiMED) and Department of Medical Sciences (DCM); Department of Chemistry, University of Aveiro, 3810-193, Aveiro, Portugal; Multidisciplinary Institute of Ageing (MIA-Portugal), University of Coimbra, 3004-504 Coimbra, Portugal

## Abstract

*Candida albicans* is a diploid pathogen known for its ability to live as a commensal fungus in healthy individuals, but causing both superficial infections and disseminated candidiasis in immunocompromised patients where it is associated with high morbidity and mortality. Its success in colonizing the human host is attributed to a wide range of virulence traits that modulate interactions between the host and the pathogen, such as optimal growth rate at 37°C, the ability to switch between yeast and hyphal forms and a remarkable genomic and phenotypic plasticity. A fascinating aspect of its biology is a prominent heterogeneous proteome that arises from frequent genomic rearrangements, high allelic variation, and high levels of amino acid misincorporations in proteins. The latter leads to increased morphological and physiological phenotypic diversity of high adaptive potential, but the scope of such protein mistranslation is poorly understood due to technical difficulties in detecting and quantifying amino acid misincorporation events in complex proteomic samples.

To address this question, we have developed and optimized mass spectrometry and bioinformatics pipelines capable of identifying low-level amino acid misincorporation events at the proteome level. We have also analysed the proteomic profile of an engineered *C. albicans* strain that exhibits high level of leucine misincorporation at protein CUG sites and employed an *in vivo* quantitative gain-of-function fluorescence reporter system to validate our MS/MS data. The data show that *C. albicans* misincorporates amino acids above the background level at protein sites of diverse codons, particularly at CUG sites, confirming our previous data on the quantification of leucine incorporation at single CUG sites of recombinant reporter proteins. The study also demonstrates that increasing misincorporation of Leucine at CUG sites does not alter the translational fidelity of the other codons. These findings advance existing knowledge on amino acid misincorporations in *C. albicans* and add a new dimension to the remarkable capacity of this fungus to diversify its proteome.

## Introduction

All living organisms face challenging environments and rely on adaptation mechanisms to survive and thrive. Pathogens that colonize various niches in the human body for instance, are constantly challenged by host immune defences, therapeutics, and variable conditions, such as temperature, pH, and nutrition (Alves *et al*., 2020). The extracellular changes activate signalling pathways that trigger transcriptional responses, leading to modulation of the fungal proteome to better respond to stimuli. While genetic mutations occur as discreate DNA events that are easy to map by sequencing, are inherited by the following generations and mutate all polypeptides encoded by the mutant protein coding genes, i.e., change the proteome and the phenome in a quantitative and heritable manner, protein biosynthesis errors produced during mRNA translation (mistranslation) affect a small percentage of the polypeptides of each protein, normally occur randomly, affect a very large number of proteins or even the entire proteome of an organism, are normally transient and not passed to the following generations. Importantly, mistranslations result in the production of statistical proteins, i.e., proteins that are a mixture of wild type and mutated polypeptides where the latter are diverse and normally represent a small percentage of the total number of polypeptides of each protein. These amino acid misincorporations have been looked as inevitable errors of the protein synthesis machinery that had no consequence or could be slightly detrimental due to their potential to destabilize protein structure in ways that could result in protein misfolding, aggregation and loss of function (Drummond & Wilke, 2009). However, the alternative hypothesis of adaptive translation, where the polypeptide variants produced by codon mistranslations increase the functional repertoire of a proteome or facilitate the rewiring or amplification of signalling networks, enhancing the organisms capacity to respond and adapt quickly to environmental changes, is gaining increasing support (Ribas de Pouplana *et al*., 2014, Mohler & Ibba, 2017).

Recent works show that mistranslation is widespread in nature and that both single cell or multicellular organisms can take advantage of it, by regulating its levels, under specific physiological and environmental conditions (Ling *et al*., 2015, Schwartz & Pan, 2017). For example, inactivation of the threonyl-tRNA synthase editing domain by oxidative stress or point mutations can lead to threonine-to-serine misincorporation in *Escherichia coli* and *Mycoplasma* spp. (Ling & Söll, 2010, Li *et al*., 2011). *E. coli* exposed to aminoglycoside antibiotics, which interfere with the ribosomés proofreading mechanism, or grown under amino acid starvation conditions mistranslates at high level (Mordret *et al*., 2019). The archaeon *Aeropyrum pernix,* which grows optimally at 90°C, undergoes inducible leucine-to-methionine substitution when incubated at 75°C, leading to enhanced enzymatic activity of hyperthermophilic proteins, likely through an increase in the conformational flexibility required for protein function (Schwartz & Pan, 2016). *Mycobacterium* species exhibit increased mistranslation in response to mutations in the GatCAB enzyme complex, increasing their resistance to rifampicin (Javid *et al*., 2014). Remarkably, GatCAB natural mutations were found in clinical isolates from *Mycobacterium tuberculosis,* providing a mechanism that enhances the microorganism’s survival during infection (Su *et al*., 2016). Mistranslation could further increase the sampling of genetic mutations by increasing tolerance to stressful agents. The artificial induction or suppression of generalized mistranslation levels affects *E. coli*’s early survival and resistance to DNA damage caused by ciprofloxacin (Samhita *et al*., 2020).

The phenomenon of mistranslation has been extensively studied in the leucine CUG codon, which has been reassigned to either serine or alanine, or ambiguously assigned to serine and leucine in several fungal species of the so called CTG clade (Krassowski *et al*., 2018). Serine-to-leucine (Ser→Leu) ambiguous decoding has been observed in *Ascoidea asiatica* (Mühlhausen *et al*., 2018), *Candida maltosa* (Suzuki *et al*., 1997), *Candida albicans* (Gomes *et al*., 2007) and more recently, in the halotolerant yeast *Debaryomyces hansenii* (Ochoa-Gutiérrez *et al*., 2022). This unique translational event creates a Leu/Ser statistical proteome i.e., a proteome where each protein is a statistical average of polypeptides containing Leu or Ser at CUG sites (Woese, 1965). The number of different polypeptides of each protein is determined by the expression n^2^, where n is the number of CUG codons in the mRNA and the power 2 refers to both Leu and Ser. In *C. albicans*, the ambiguous decoding of CUG codons allows for the potential generation of >1.0 ×10^11^ different polypeptides from its 6226 genes (Gomes *et al*., 2007).

The impact of CUG dual translation has been thoroughly analysed in *C. albicans,* particularly in the context of the commensal-pathogen transition, host-pathogen interactions, and immune evasion. Accumulating evidence suggests that the identity of the CUG codon can rewire protein-protein interactions (Stynen *et al*., 2010, Côte *et al*., 2011) and modulate protein activity in important virulence traits (Rocha *et al*., 2011, Sárkány *et al*., 2014). These CUG encoded residues are often located in conserved regions of proteins involved in biofilm formation, mating, morphogenesis, adhesion, and signal transduction. For instance, the substitution of polar serine by non-polar leucine in *C. albicans* leads to a decrease in the stability and activity of the Cek1 signalling kinase (Fraga *et al*., 2019), an increased substrate adherence by cells expressing the adhesin Als3-Leu (Miranda *et al*., 2013) and to temperature sensitivity of the translation initiation factor (eIF)4E, which mediates mRNA binding to the ribosome (Feketová *et al*., 2010). Despite a high fitness cost in rich medium, *C. albicans* tolerates high level of leucine incorporation at CUG sites (Bezerra *et al*., 2013) and hypermistranslating strains are more resistant to oxidative stress and antifungals, display high phenotypic diversity (Miranda *et al*., 2007), have higher adherence to substrates, and are less susceptible to phagocytosis (Miranda *et al*., 2013). These observations are relevant given the interaction of *C. albicans* with its human host.

In *C. albicans,* CUG translation is facilitated by a hybrid tRNA(CAG)Ser that contains identity elements for both seryl-tRNA synthetase (SerRS) and leucyl-tRNA synthetase (LeuRS). This results in the aminoacylation of the Ser CUG decoding tRNA with Ser (Ser-tRNA(CAG)Ser) or Leu (Leu-tRNA(CAG)Ser) and consequent incorporation of Ser or Leu at CUG codons at the ribosome A-site. The incorporation of Leu at CUG sites is possible because the mischarged Leu-tRNA(CAG)Ser is not edited by the LeuRS editing site and is not discriminated by the translation elongation factor1 (eEF1A) (Santos *et al*., 1996). Under normal physiological conditions the tRNA(CAG)Ser is mainly aminoacylated with Ser by the SerRS (appx 97%) (Gomes *et al*., 2007).

Different reporter systems have been employed to quantify CUG mistranslation. One approach relies on the GFP marker whose stability and activity depend on Leu incorporation at codon site 201 (Bezerra *et al*., 2013). Incorporation of Ser at position 201 leads to degradation and full inactivation of GFP. However, introduction of CUG_201 by site directed mutagenesis leads to partial recovery of fluorescence which can be correlated with the levels of mistranslation (Leu misincorporation) by microscopy or flow cytometry. Another method involves mass spectrometry (MS) of a specific protein containing a single CUG site. The use of synthetic peptides for the calibration of the MS system enables the quantification of Ser and Leu peptide variants (Gomes *et al*., 2007). However, these approaches are limited to single CUG sites and accurate data on Leu and Ser incorporation on a proteome wide scale has been difficult to obtain. This is due to a lack of MS sensitivity to detect the rare peptides containing amino acid misincorporations that normally appear in complex protein samples, normally below 0.1% of the wild type peptides, and to statistical and computational difficulties in dealing with the high noise-to-signal ratio of the MS datasets. However, recent works provide some evidence that these difficulties could be overcome and raise the hope that it may be possible to detect rare peptide variants in complex mixtures of peptides produced from total protein samples and identify isoforms that are absent from reference protein databases (Dupree *et al*., 2020). In this study, we developed a pipeline based on PEAKS software algorithms (Bioinformatics Solutions Inc.) (Han *et al*., 2005) to detect and determine the frequency of low level amino acid misincorporations in complex peptide mixtures prepared from total protein extracts of *C. albicans*.

The study demonstrates that our MS-based method is sufficiently robust to identify rare mutant peptides and determine translation error frequencies in complex peptide mixtures produced from total protein extracts of *C. albicans*. We have validated our MS data using bioengineered strains of *C. albicans* whose leucine misincorporation at serine CUG sites was 20%, as determined using our GFP gain of function reporter system and single protein-single site MS analysis (Gomes *et al*., 2007, Bezerra *et al*., 2013). The data show that CUG is not the only ambiguous codon in *C. albicans*, and that the engineering of high levels of leucine misincorporation at CUG sites does not alter the translational accuracy of other codons. Our findings also indicate that further developments in protein sample preparation, protein fractionation, MS sensitivity and computational data analysis are required to overcome the high noise-to-signal ratio associated with rare peptides and accurately quantify translational amino acid misincorporations at the global proteome level. Finally, this work provides a promising opportunity to detect and determine the frequency of protein biosynthesis errors in different wild-type clinical strains of *C. albicans* and of other pathogens, as well as in other cell types, leading to a better understanding of the role of translational errors in microbial infections and diseases in general.

## Results

### Tools to detect amino acid misincorporations in *Candida albicans*

Protein biosynthesis errors occur frequently, with estimated rates of 10^-5^ to 10^-4^ errors per incorporated amino acid in yeast (Kramer *et al*., 2010), which is much higher than the mutation rates of DNA replication (10^-9^ to 10^-10^) (Lynch *et al*., 2008, Zhu *et al*., 2014) and transcription (10^-6^) (Chung *et al*., 2023). Translational errors at frequencies of 1 misincorporation in 10,000 to 100,000 codons translated by the ribosome produce extremely low abundance peptides whose detection and identification are highly challenging. To overcome such limitations, misincorporations have been detected at single preselected amino acid sites by monitoring misincorporation of radioactive amino acids or by the recovery of function of mutant fluorescent or chemiluminescent reporter systems, where gain of function depends on misincorporation of wild type amino acid at the preselected sites (Tavares *et al*., 2018).These methods are sensitive and accurate but are restricted to single codon positions of specific reporter proteins and do not provide a global view of amino acids misincorporations, which is critical to fully understand the biology of protein biosynthesis errors. Recent works carried out by our group and others have attempted to resolve this issue by analysing amino acid misincorporations in highly complex mixtures of total protein extracts from bacteria, yeast, and mammalian cells, by mass spectrometry (MS) (Mohler & Ibba, 2017, Mordret *et al*., 2019, Varanda *et al*., 2020). These studies produced promising data but also highlighted the need to develop new sample preparation and peptide fractionation methods as well as more robust computational methods to search the MS data space. In this study, we present a pipeline for the identification of amino acid misincorporations and for determination of codon-associated error frequency at a global proteomic scale in the pathogenic fungus *C. albicans* mass spectrometry (MS).

*C. albicans* total protein extracts were prepared using current methods and proteins were fractionated by gel electrophoresis into 8 fractions. Proteins were digested with trypsin and the resulting peptide mixtures of each fraction were subjected to downstream LC-MS/MS analysis. Data analysis was carried out using the PEAKS DB algorithm, which integrates database search with *de novo* sequencing, to identify peptides and proteins. The haploid complement of features of the diploid Genome Assembly 22 available at the Candida Genome Database (CGD) was used as a reference database for this analysis.

To evaluate the quality and representativeness of the samples, we assigned each amino acid present in the final list of identified peptides to its corresponding codon and determined its overall frequency in the sample. The frequency pattern of abundant/rare codons and the relative synonymous codon usage (RSCU) of the wild-type sample (T0 strain) were similar to those of the reference database (Figures 1 and S1), indicating that the detected peptides accurately represented the global codon usage. However, some deviations were observed in rare codons. For example, 14 out of the 16 codons with a frequency in the reference genome below 6.5 codons per thousand (half of the codonś frequency median) were underrepresented in the T0 strain sample, and the most underrepresented codons were rare codons. We then focused our analysis on the CUG codon, which is associated with ambiguous Ser/Leu translation in *C. albicans*. The CUG codon has a genomic frequency of 4.2 per thousand codons (0.33 RSCU) but the data of our samples showed a lower frequency of 1.8 per thousand codons (0.17 RSCU), i.e., the MS approach used in this work misses peptides containing rarely used codons. Indeed, proteins encoded by genes containing rare codons are normally expressed at low level, as indicated by the codon adaptation index – CAI (Gomes *et al*., 2007). Reassuringly, the peptides that were predominantly detected corresponded to abundant proteins that contained few CUG codons (average of 0.14 CUGs per proteins with area >1.0×10^8^) while peptides of proteins with low associated total areas, where the average of CUG frequency was 3.15, were rarely detected (Figures 2A and 2B). In the specific case of the CUG peptides detection, sensitivity may have also been affected by a highly biased accumulation of these codons in genes encoding membrane and cell wall proteins, whose hydrophobicity and low solubility decreases their abundance in total protein extracts (Rocha *et al*., 2011, Kachuk *et al*., 2016). Accordingly, we observed an underrepresentation of membrane proteins in our MS sample, which contained fewer CUG proteins than the reference genomic database and a positive bias towards proteins containing a lower number of CUG codons (Figure 2C and 2D). These technical issues likely decreased the ability to detect amino acid substitutions at rare codon sites in our samples. However, we were still able to identify 54% of the *C. albicans* proteome, with most proteins (>60%) identified by more than 4 peptides and an average amino acid sequence coverage of 31% (Figure S2). We used this raw dataset to analyse the profile of amino acid substitutions in *C. albicans* by uncovering peptide variants with mass shifts relative to the genome-encoded sequences.

**Figure 1.**
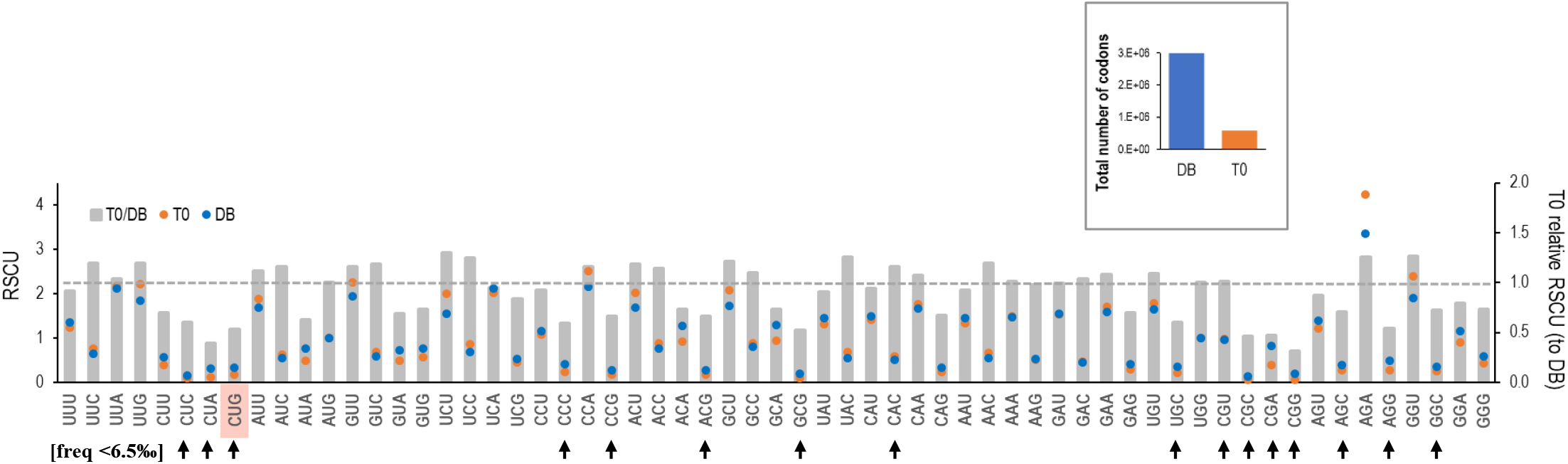
Codon frequency analysis. Dots in the graphic compare the relative synonymous codon usage (RSCU) for the reference proteome used for PEAKS DB search and for the T0 strain sample after codon assignment to all identified peptides (without duplicates). The columns highlight the RSCU deviations of the T0 strain sample regarding the RSCU obtained for the reference DB, showing their RSCU ratio. The dashed line indicates the expected ratio if no differences were observed. Arrows point to rare codons, defined as those whose frequency is lower than half of the median of all codon frequencies obtained for the DB used for reference and calculated by Anaconda software (6.5 codons per thousand). The box shows the total number of codons used for this analysis.

**Figure 2.**
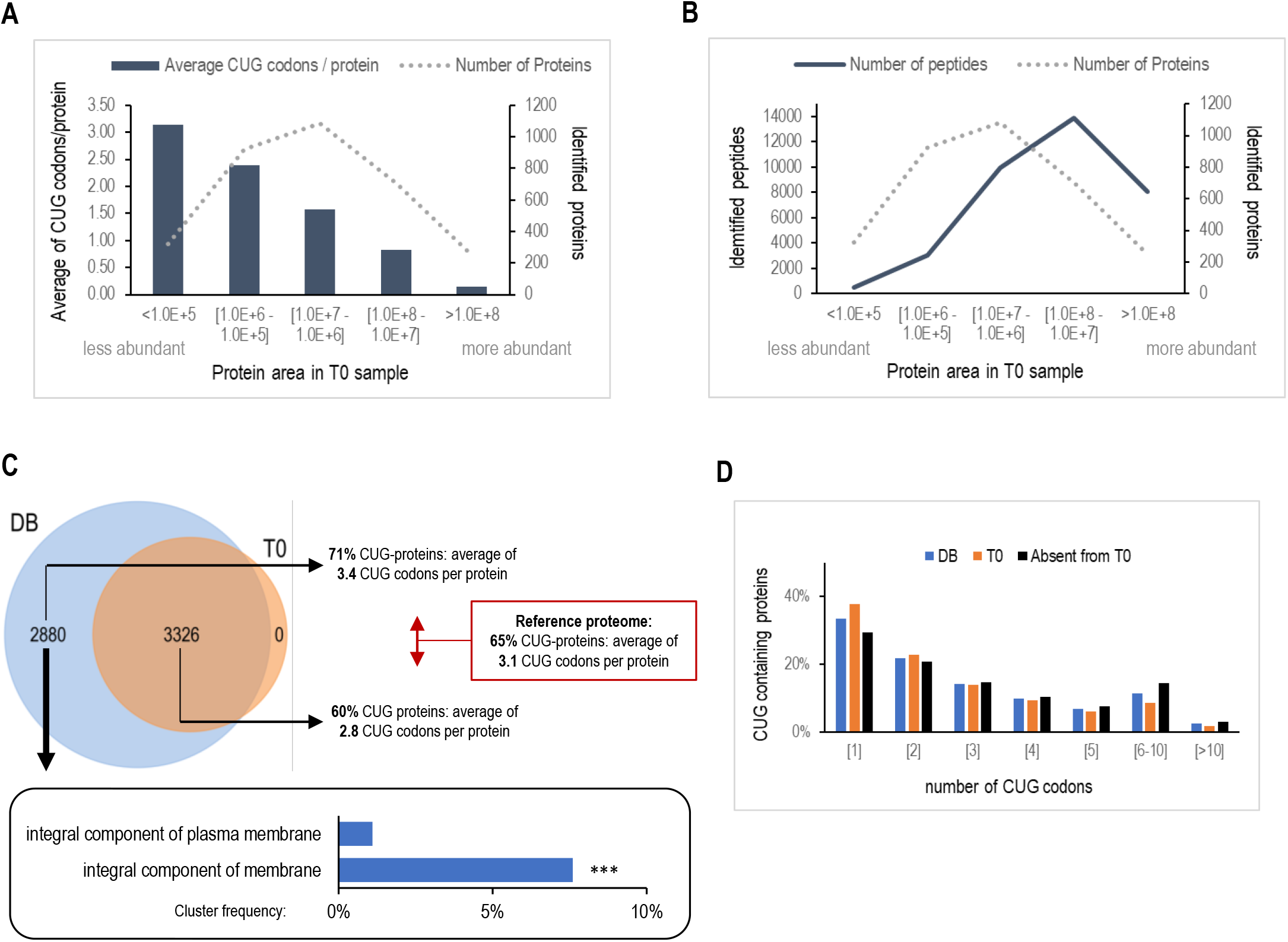
CUG codons are underrepresented in the peptides of wild-type MS sample. **A)** Link between protein abundance and CUG content. **B)** Link between protein abundance and number of identified peptides. **C)** Venn diagram and gene ontology of unidentified proteins. Proteins in both fractions have been analysed regarding their CUG content. Venn tool available in the program FunRich 3.1.3. GO enrichment analysis of proteins absent from the MS sample, using GOTermFinder application at CGD (background: 6206 proteins from MS reference database). p-value cut-off: 0.05. **p<0.01; ***p<0.001. **D)** Analysis of identified proteins regarding their content on CUG codons.

An additional difficulty was related to the diploid and heterozygous nature of the *C. albicans* genome, which complicated the differentiation of translational errors from alternative amino acid incorporations associated with genomic allelic variations. To circumvent this issue, we used the diploid reference proteome available at CGD (Figure S3), which contains the translated sequences of both haplotypes (allele A and allele B), as a reference database for peptide-spectrum matches. To reduce redundancy and maintain sensitivity, we removed from the *C. albicans* diploid database all sequences from allele B when allele A of the same protein had an identical amino acid sequence. For spectra that did not match the pre-selected database during the *de novo*-assisted database search, the PEAKS software contains algorithms that detect post-translational modifications (PEAKS PTM) and mutations (SPIDER). We used the pipeline described in Figure 3, assuming that mutated peptides (mistranslated) are less abundant than their unmutated counterparts and that post-translational modifications (PTMs) with isobaric delta masses can better explain peptide mass shifts than amino acid misincorporations.

**Figure 3.**
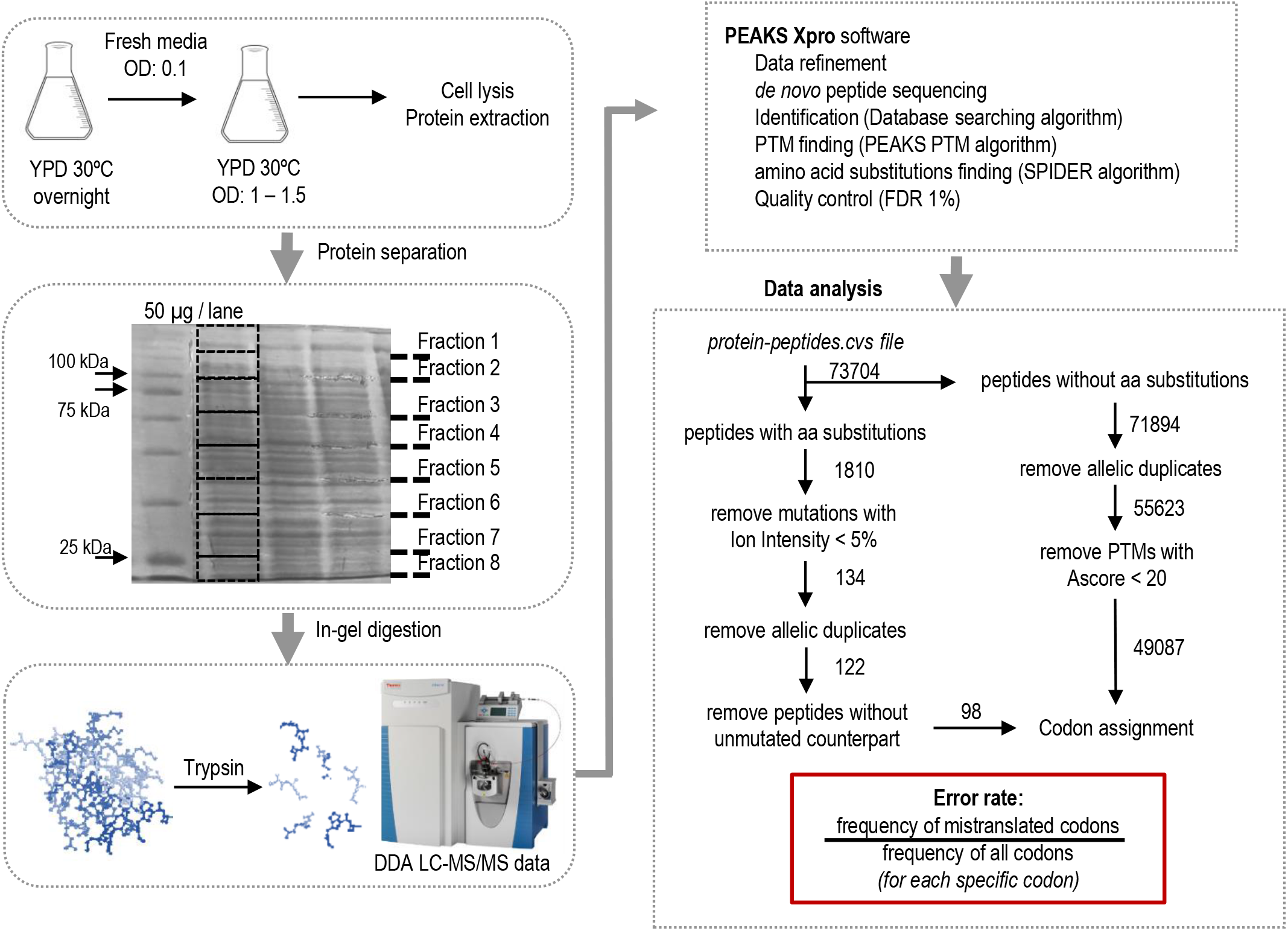
Workflow for mistranslation analysis. Protein extracts from actively growing cultures were resolved in SDS-PAGE and divided into 8 fractions. Bands were manually excised from the gel, digested, and injected separately in the MS system. Raw MS/MS data was analysed by PEAKS Xpro software. The list of identified peptides was filtered to remove duplicates and low-quality hits, and codon assignment and mistranslation frequency was calculated using R scripts. Numbers indicate the number of peptides identified or validated in each step of the pipeline from T0 (wild-type) strain.

We analysed and filtered the list of all identified peptides (*protein-peptides* file – Supplementary data) from a *C. albicans* wild-type strain (T0 strain) using R scripts (see Materials and Methods). Peptides with low-quality PTMs that did not pass the AScore filter were removed from the list, as well as peptide duplicates with the same amino acid sequence, matched to the same protein, but to a different allele. For correct codon-amino acid assignments we took into consideration that peptides of identical sequences could be aligned to different proteins due to the presence of paralogs and conserved regions among protein families. To minimize overestimation of codon frequencies, a single protein per peptide was chosen for codon assignment and we also developed our scripts to record any potential bias in codon assignment. Additionally, for peptides that aligned to multiple regions of a protein due to the presence of repetitive regions the program assumed the position closer to the N-terminal of the protein. Finally, to validate peptides with amino acid misincorporations, we used an ion intensity filter to localize the mutated amino acid (≥5%) with high accuracy and accepted only those peptides whose unmutated counterpart (wild type peptide, correctly translated) was also detected in the sample.

### Protein biosynthesis errors occur at different codons

To evaluate global mistranslation, we utilized the SPIDER algorithm of the PEAKS software, which reconstructs new peptide variant sequences by combining homology search and *de novo* sequence tags (Han *et al*., 2005). After applying database search and validation filters, the algorithm identified several amino acid substitutions in different codons of 540 peptides. Before assuming these variants as true mutations we have considered the possibility that PTMs and mutations could produce identical mass shifts and that PTMs are more frequent than amino acid misincorporations (Kim *et al*., 2016). For instance, a mass shift of 14.02 Da can result from either methylation or Gly to Ala, Asp to Glu, Val to Ile/Leu, Ser to Thr or Asn to Gln substitutions. To avoid inaccurate assignment of amino acid misincorporations, which would overestimate mistranslation events, we excluded any substitution whose associated mass shift could be explained by a PTM. For this, the PEAKS PTM identification algorithm was applied to the data upstream of the SPIDER algorithm and as expected, the number of misincorporations decreased (Figure 4). By following this workflow, and after removing redundant peptides due to multiple protein alignments, 94 unique mutated peptides were detected out of 48474 peptides identified in the wild-type strain (T0). These mutations occurred in 27 different codons (44 substitution types), and the most frequent substitutions detected were alanine-to-glutamine (Ala_GCU/GCC_→Gln) and glycine-to-asparagine (Gly_GGU_→Asn). We also observed that serine codons were prone to aspartic acid misincorporation, although serine formylation can occur as an artefact (Lenčo *et al*., 2020), resulting in the same mass shift as Ser→Asp substitution.

**Figure 4.**
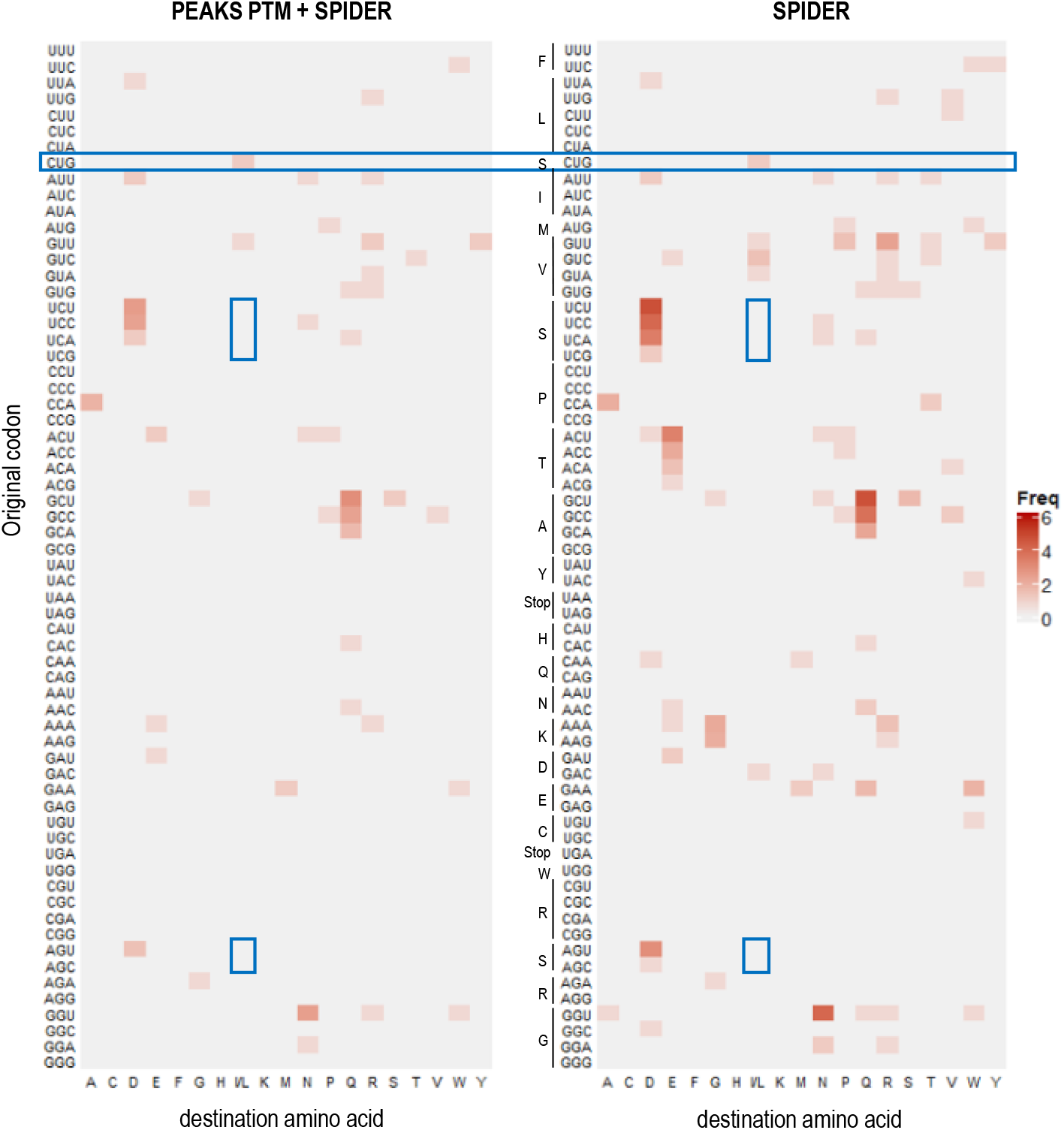
Impact of using PEAKS PTM algorithm before SPIDER in the identification of amino acid substitutions. The matrix reveals the number and identity of substitutions found for each codon. Different post-translational modifications with isobaric delta masses may explain the observed differences. Highlighted in blue boxes are amino acid substitutions in the CUG codon and Ser→Leu substitutions in serine codons.

In line with previous reports (Akashi, 1994, Sun & Zhang, 2022), we observed that the codons prone to mistranslation were not necessarily the most frequently used in the peptides of our sample. Indeed, highly frequent codons like GAU, AAU, and CAA showed high translational accuracy, indicating that biosynthesis errors are not correlated with codon usage frequency (Figure S4). As expected, we were only able to detect two mistranslation events in rare codons, specifically His_CAC_→Gln and Ser_CUG_→Leu. The Ser/Leu ambiguous translation of CUG codons in *C. albicans* has been previously described and quantified using fluorescent reporters and MS/MS of recombinant reporter proteins (Gomes *et al*., 2007, Bezerra *et al*., 2013). Our new data confirmed this mistranslation event, and we observed no other substitutions at the CUG codon, even when PEAKS PTM was not used. Furthermore, the data indicates that this substitution only occurs at the CUG codon and not at any other of the six serine codons (Figure 4).

### The CUG codon is mistranslated at high frequency in *C. albicans*

To gain a comprehensive understanding of the distribution of translational errors in the *C. albicans* proteome, each amino acid of the identified and validated peptides was assigned to the corresponding codon. Translational accuracy was assessed by calculating error frequency, defined as the number of times a specific codon is mistranslated relative to the total number of times it appears in the MS dataset. The global error frequency for the wild-type sample (T0 strain) was 1.08 x 10^-4^ (0.01%), considering the median of all codon-associated mistranslation frequencies (Figure 5A). While GCC, GCU, and GGU codons were found to be associated with the most frequent amino acid misincorporations, their calculated error frequency of 0.08%, 0.06%, and 0.03%, respectively, was lower than the error frequency for the rare CUG codon (0.17%). We detected more than one amino acid misincorporated at both GCC and GCU codon sites, but Ser→Leu were the only substitutions detected at CUG sites (Figure 5B). Thus, the CUG codon had the highest mistranslation frequency of all amino acid-codon pairs, (Figure 5A, left side) and this result was maintained even when the error frequency associated with each codon independently of the misincorporated amino acid was computed (Figure 5A, right side).

**Figure 5.**
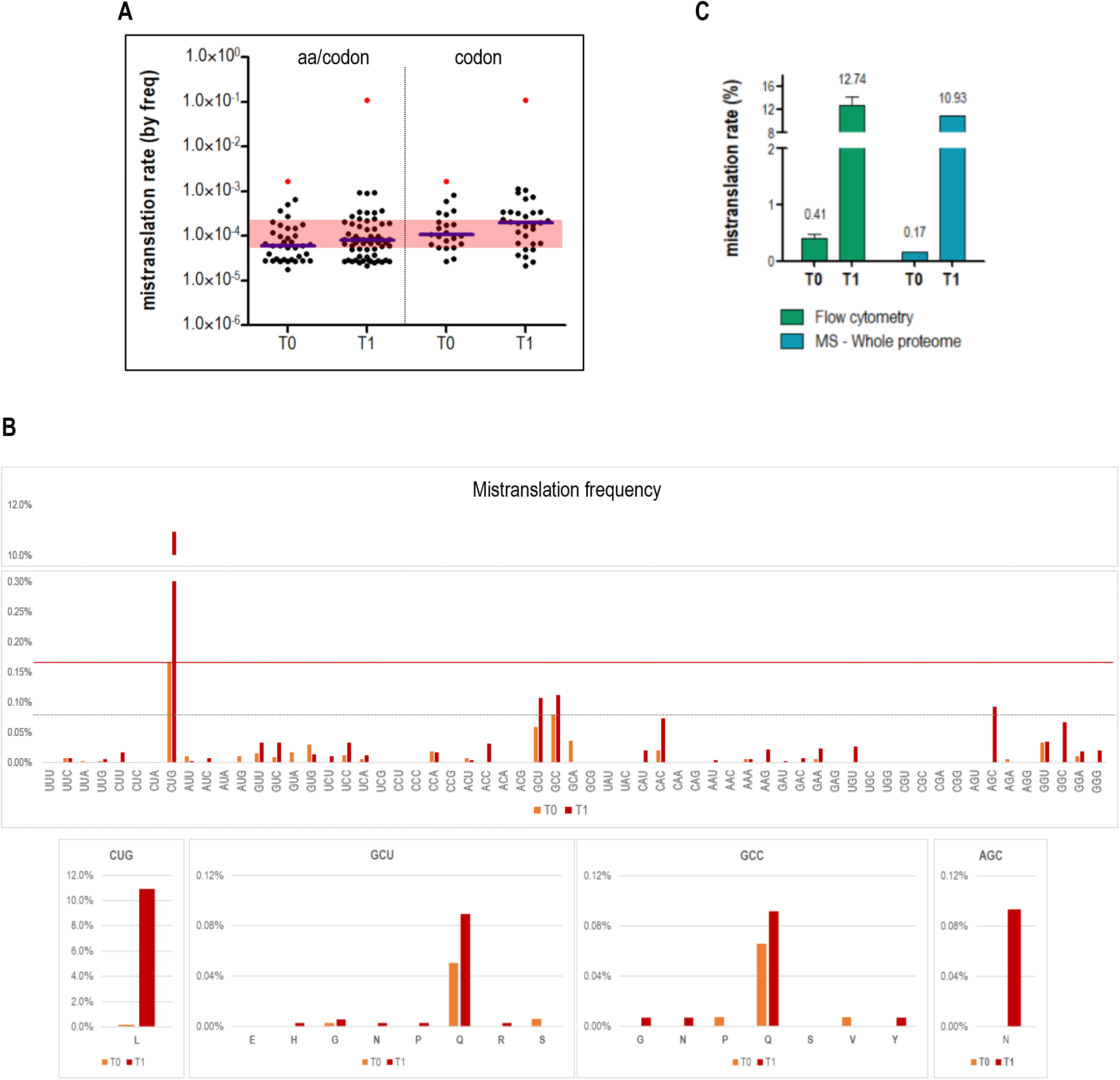
Comparative analysis of protein biosynthesis errors in strains with different levels of Leu misincorporation at CUG codons. Mistranslation frequency was assessed by LC-MS/MS analysis of whole cell lysates **(A and B)** and by flow cytometry to detect a fluorescent reporter **(C)**. Error bars from flow cytometry data represent the standard deviation of the mean from three independent experiments. **A)** Distribution of mistranslation frequency calculated for each codon mutation either specific for one single amino-acid (left) or for all detected substitutions (right). The blue line indicates the median of all calculated mistranslation frequencies. CUG codon mistranslation frequency is highlighted in red. **B)** Comparison between codons mistranslation frequencies of T0 and T1 strains. Graphics at the bottom detail which amino acids are being misincorporated and at what frequency in codons whose mistranslation frequency is closer to the one found for CUG in the wild-type strain (>0.08%).

Previous studies have quantified CUG Ser/Leu ambiguity in *C. albicans* using mass spectrometry analysis of purified recombinant proteins (Gomes *et al*., 2007) and fluorescence microscopy of strains transformed with fluorescent reporters (Bezerra *et al*., 2013). The error frequency estimated by our pipeline was lower than the error rate obtained by microscopy (1.45 ± 0.85%) and direct MS analysis (2.96 ± 0.49%) at single sites of a purified protein. However, it was similar to that obtained by flow cytometry (0.41 ± 0.06%) where a very large number of cells are monitored and the standard deviation of the data is lower (Figure 5C), demonstrating the reliability and effectiveness of our method. Moreover, to validate our pipeline, we analysed a Leu-CUG hypermistranslating strain (T1) engineered in our laboratory which misincorporates 20.61 ± 1.81% of Leu at the Ser-CUG sites, as quantified by fluorescence microscopy (Bezerra *et al*., 2013). Consistent with the higher level of CUG-Leu incorporation, we detected 98 Ser→Leu substitutions in peptides derived from 84 proteins of the T1 strain, contrasting with only 2 peptides in 2 proteins of the T0 control strain. Importantly, almost all CUG-related Leu misincorporations in the T1 strain occurred in proteins of the T0 strain where we could detect the wild-type CUG peptides (Ser-CUG incorporation only). This provides a clear indication that proteome scale detection of rare peptides containing amino acid misincorporations requires the development of more sensitive sample preparation, MS/MS, and computational methods.

To rule out the remote possibility that misincorporations at CUG sites resulted from genetic variation related to genetic manipulation of the T0 and T1 strains, we reanalysed the genome sequencing data of these strains (Bezerra et al., 2013), focusing on the genes that encoded the peptides containing amino acid misincorporations in both strains. No allelic difference was detected at those specific sites (Table S1), confirming that the mutations were true translational amino acid misincorporations. Moreover, a stringent analysis of putative error arising from codon misassignment related to alignment of peptides to more than one protein was also carried out. Four mutated peptides matched different proteins in the T0 strain, but only one mutated peptide (K.NQQ(sub A)AMNPANTVFDQ(sub A)K.R) with equal area, mass and retention time, matched with the same probability proteins that are genetically distinct, C1_13480W_A and C1_04300C_A. The second Q(sub A) could be attributed to either GCC or GCU codons, leading to potential alterations in codon frequencies. In the T1 strain, eight mutated peptides with different protein matches were identified, four of them at genetically different chromosomal loci: two involving GCC/GCU mismatches (with no change in the error frequency), and the other two peptides whose mismatches were assigned to CAC codons instead of CAU codons, leading to a change in the codon error frequency from 0.07% and 0.02%, respectively, to 0.04% for both. None of the identified CUG-containing peptides in either T0 or T1, were matched to more than one protein. The estimated CUG mistranslation frequency in the T1 strain (10.94%) was lower than that reported previously, but similar to the *in vivo* values obtained by flow cytometry (12.74 ± 1.20%). By using this pipeline, we were thus able to discriminate two *C. albicans* strains with different CUG-related error frequencies.

### The artificial increase of leucine incorporation is CUG-specific

*C. albicans* CUG-hypermistranslating strains exhibit a high degree of phenotypic and genomic diversity, which impacts fungal adhesion to host substrates, immune recognition, and tolerance to antifungals (Bezerra *et al*., 2021). However, it is unclear whether these adaptive phenotypes result from Ser/Leu substitutions at CUG sites or from general deregulation of translational fidelity caused by the stress induced by Ser→Leu CUG mistranslation and/or by the heterologous expression of the *Saccharomyces cerevisae* tRNA(CAG)Leu and the GFP reporter. Our new data show a 3-fold increase in the number of mutated peptides in the T1 strain relative to the wild-type T0 strain (in a total sample of 48474 peptides in T0 versus 50186 peptides in T1), with an overall mistranslation frequency of 1.95×10^-4^ (0.02%) in T1 versus 1.08 x 10^-4^ (0.01%) in T0 (Figure 5A). The analysis of the identity and frequency of substitutions confirms a bias towards mistranslation of CUG codons and the specificity of Leu incorporation at CUG sites. Indeed, Leu was not incorporated at any other of the six Ser codons (Figure 5B and S5), and the translational accuracy of other codons was not affected. Therefore, these data confirm for the first time that the phenotypes previously observed in the T1 strain (Bezerra *et al*., 2013, Weil *et al*., 2017) are a direct result of increased Leu/Ser ambiguity at CUG sites.

Interestingly, the total number of CUG codons assigned to the T1 peptides was lower relative to the number detected in the control T0 strain (897 in T1 vs 1197 in T0). This, combined with the higher number of mistranslated codons (98 in T1 vs 2 in T0) explained the increase in misincorporation frequency from 0.17% in T0 to 10.93% in T1. To further clarify the result above, we estimated the RSCU in T0 and T1 strains using the detected peptides as described in the methods section. Unlike other codons that showed similar RSCU values in both strains, CUG codons were underrepresented in T1 (Figure 6A) suggesting that Leu misincorporation at certain CUG sites may lead to protein misfolding and degradation, as previously analysed in our laboratory (Rocha *et al*., 2011). However, when analysing the RSCU using the full sequence of all identified proteins, no differences were observed. The percentage of identified proteins with CUG-encoded residues was similar between T0 (60.5% with an average of 2.81 CUG codons per protein) and T1 (59.7%, average of 2.84). The major difference was the detection of peptides containing CUG-encoded residues, which were identified in 34.7% of the T0 proteins and only in 24.0% of the T1 proteins. Apart from this, there were no major differences found regarding proteome coverage, overall protein sequence coverage and the number of identified peptides per protein (Figure S2). In line with this, the higher level of Leu incorporation at CUG sites in the T1 strain remodelled the proteome: 263 and 354 proteins were exclusive of T0 and T1, respectively. A GO enrichment analysis showed that T0-specific proteins were associated with ubiquitination and autophagy, and more than 80% of these proteins contained at least one CUG codon (Figure 6B and Figure S6A). Interestingly, the function of 40% of the T1-specific proteins is “unknown”. We used the PEAKS Q module from PEAKS Xpro software to perform a label-free quantification on the proteins common to both strains and found that 680 were down-regulated and 247 were up-regulated in the hypermistranslating strain relative to the T0 control (Figure 6C). Although there were only slight differences in the total abundance of common proteins (Figure 6D), we found that 62% of the down-regulated proteins contained CUG codons and were involved in regulating transcription by RNA polymerase I (Figure S6B). On the other hand, the up-regulated proteins participated in translation, peptide, amide, and pyrimidine biosynthetic and metabolic processes, and contributed to the structural integrity of the ribosome.

**Figure 6.**
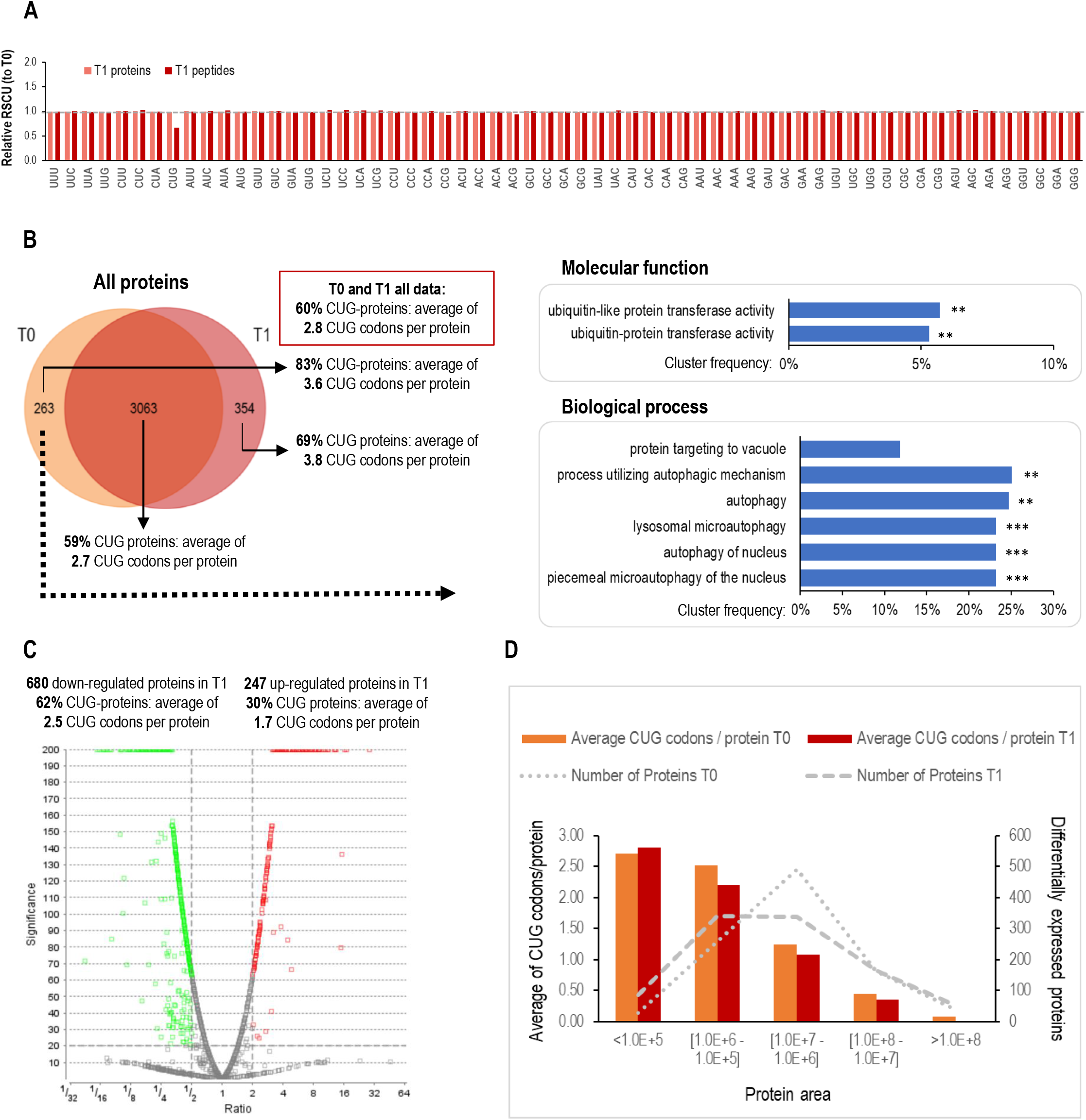
Comparative proteogenomic analysis between T0 and T1 strains. **A)** Codon frequency analysis. RSCU deviations of the T1 sample regarding the RSCU obtained for the wild-type (T0) sample, calculated from all detected peptides or from all identified proteins using Anaconda software. The dashed line indicates the expected ratio if no differences were observed. **B)** Venn diagram and gene ontology of T0-specific proteins. Common and exclusive proteins identified in T0 and T1 strains were analysed regarding their CUG content. Venn tool available in the program FunRich 3.1.3. GO enrichment analysis of proteins absent from T1 sample, using GOTermFinder application at CGD (background: 3680 proteins identified in both strains). p-value cut-off: 0.05. **p<0.01; ***p<0.001. **C)** Volcano plot from label free quantification of T1 relative to T0 strain obtained by PEAKS Q module. Markers for the proteins that are above the set significance threshold are displayed in red (for up-regulated proteins regarding the reference strain T0) and green (for down-regulated proteins). Up and down-regulated proteins were separately analysed regarding their CUG content. **D)** Comparison between the abundance of differentially expressed proteins and their content on CUG codons.

### Peptide exclusion lists as a strategy to increase the detection of mistranslated peptides

To optimize our methodology and increase the detection of mistranslated peptides, we tested different approaches to acquire and analyse the MS/MS data. We lessened the stringency of our data analysis filters, namely by using a mass tolerance of 10 ppm instead of 5 ppm for precursor ions during searches, allowing a semi-specific cleavage instead of trypsin-specific cleavage, and using a 1% ion intensity filter instead of 5% to validate amino acid substitutions. These variables increased slightly the number of detected peptides containing mutations but did not substantially alter the mistranslation frequencies, and the CUG codon remained the most ambiguous codon in *C. albicans* (Figure S7A).

We also investigated whether a more direct search for Leu misincorporation, by selecting Ser→Leu substitutions as a variable PTM when performing the DB search, would increase the number of peptides with Leu/Ser mutations. This strategy resulted in the identification of an additional CUG site mutation in the wild-type sample and four mutations associated to other, more frequent, serine codons. Out of the 76 new Ser→Leu mutations detected in the T1 strain, 70 were found to occur at CUG sites, suggesting a higher propensity for Leu misincorporation at CUG codons in this strain. The error frequency linked to the CUG codon was noticeably higher than that of other serine codons in both strains when employing this approach (Figure S7B).

To increase the sensitivity of our MS/MS methodology, we investigated the impact of using exclusion lists between additional runs of MS/MS acquisition, which would enable the detection of other less abundant peptides. When the samples were reinjected for a second analysis, excluding from MS/MS analysis the most abundant spectra/peptides already analysed in the first MS/MS run, we identified 343 new proteins, detected almost 13,500 new peptides, and found 29 more substitution types (Figure 7A and B). The global mistranslation frequency did not change considerably, decreasing from 1.08 x 10^-4^ (0.011%) to 9.73 x 10^-5^ (0.010%). However, there was a substantial increase in alanine to glutamine (Ala_GCU/GCC_→Gln) substitutions, which was the most frequent substitution detected with only one MS analysis.

**Figure 7.**
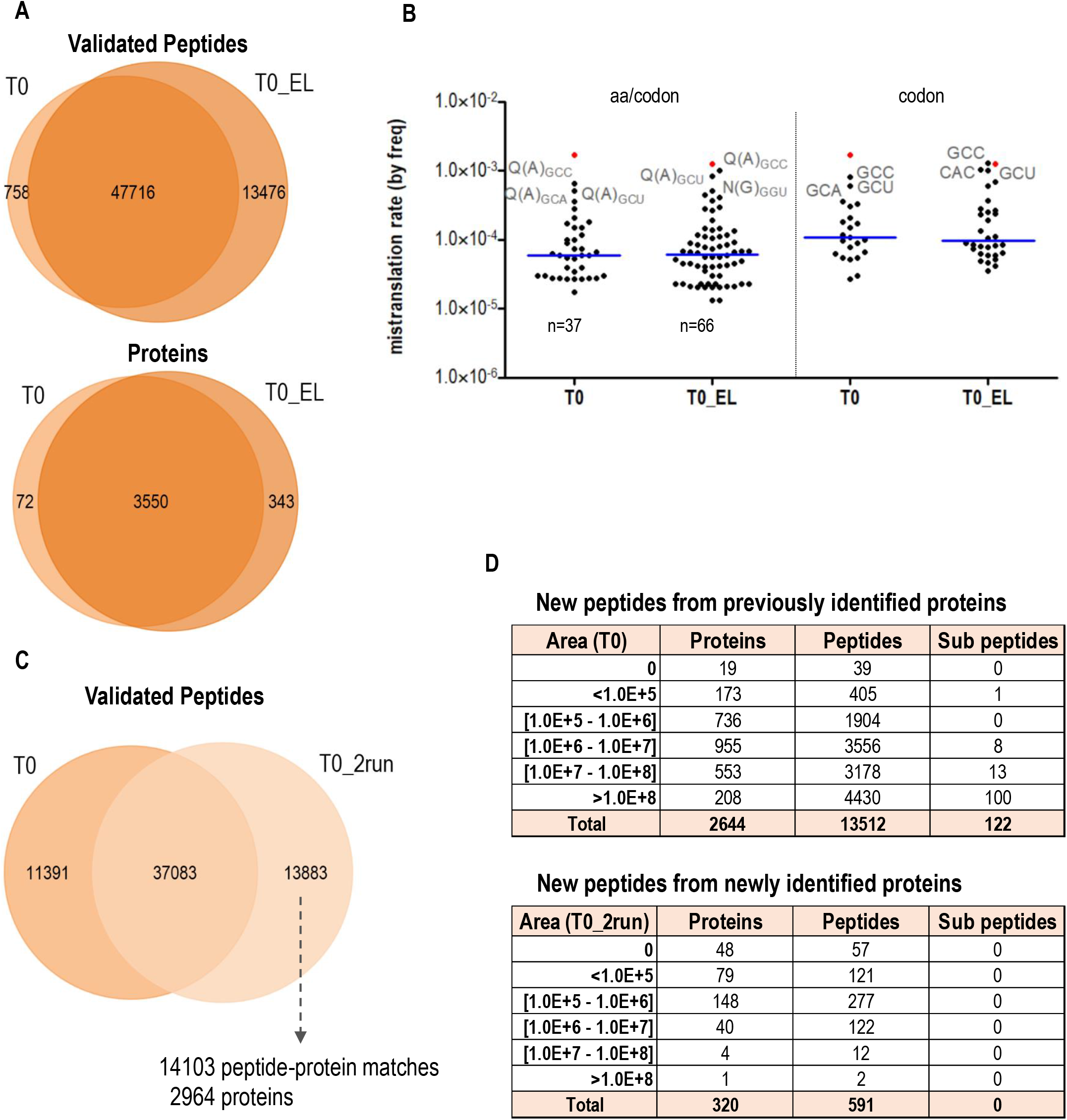
Impact of exclusion lists in the detection of peptide variants. **A)** Venn diagram showing the number of proteins and peptides identified and validated when doing the analysis with the data obtained only in the first MS injection (T0) and when using the data from both MS injections, before and after applying the exclusion lists (T0_EL). **B)** Distribution of mistranslation frequency calculated for each codon mutation either specific for one single amino-acid (left) or for all detected substitutions (right). “n” denotes the number of different substitution types and the blue line indicates the median of all calculated mistranslation frequencies. L(S)CUG mistranslation frequency is highlighted in red and the top-3 codons with highest error frequency are labelled. **C)** Venn diagram showing the number of peptides identified and validated in the first MS injection (T0) and in the second MS injection after applying the exclusion lists (T0_2run). **D)** Detailed information on peptides exclusively found in T0_2run data, correlating the protein abundance with the number of identified proteins, peptides, and peptides with amino acid substitutions.

The second MS/MS injection alone enabled the detection of 13883 new peptides (Figure 7C). We observed that the new peptides were mostly from highly abundant proteins that were previously identified in the first MS/MS analysis, and that only 4% of these new peptides led to the identification of new proteins (Figure 7D), suggesting that exclusion lists can improve peptide and protein identification sensitivity in MS analysis of complex samples, but do not substantially increase the number of peptides belonging to less abundant proteins. In fact, while the second MS/MS injection increased the detection of peptides with amino acid substitutions, they were all matched to proteins already identified without exclusion lists and mainly to those most abundant in the sample (area > 1 x 10^8^). Moreover, these new substitutions were found in codons previously identified as the most frequently associated with mistranslation such as GCU, GGU and GCC. In summary, while this is a promising strategy to increase the identification of peptide variants, it may introduce bias towards mistranslation of highly abundant proteins (and frequent codons) that could compromise our approach for global error frequency calculation, while considerably increasing the cost of the assay.

### Analysis of CUG-associated misincorporations in publicly available MS data repositories

We utilized our bioinformatics workflow to scrutinize the protein MS raw data of *C. albicans* that have been deposited in the Proteomics Identifications database (PRIDE public data repository). Over 80 samples (including replicates) from 9 different projects were reanalysed and Leu-CUG misincorporations were detected in 18 samples from 3 projects (Table S2). This limited Leu-CUG detection could be attributed to the different methodologies employed in sample preparation, fractionation, and in acquiring the MS datasets, which are optimized for each study’s objectives, and highlights the technical difficulties in analysing amino acid misincorporations by MS/MS. Indeed, most MS-based studies focus on identifying and quantifying differentially expressed proteins in distinct strains, morphologies, or under diverse environmental conditions where the noise-to-signal ratio is normally low, ignoring the presence of peptide variants from protein isoforms that are rare in complex samples.

We identified a single peptide containing a CUG Ser->Leu substitution in six samples out of 52 samples (PXD020195), which represented several morphologies and culture conditions. These samples were obtained to generate a *C. albicans* spectral library for Data Independent Acquisition (Amador-García *et al*., 2021). We also detected this amino acid misincorporation in three replicates from a dataset (PXD031774) obtained from a mixture of five isogenic strains retrieved from a fluconazole treated AIDS patient who suffered from recurrent oropharyngeal candidiasis (Song *et al*., 2022). Finally, we re-analysed a dataset (PXD027278) containing MS raw data from various *C. albicans* strains and growth forms. This dataset was originally produced to assess CUG mistranslation in various species and growth forms (Mühlhausen *et al*., 2021). In this work, Mühlhausen and colleagues refute the idea that *C. albicans* misincorporates Leu at CUG sites and argue against its prevalence as a mistranslation event in the cell. However, by using our bioinformatics pipeline, we have identified Leu-CUG incorporation events in several samples. Amino acid substitutions were also observed in other codons, but the frequency of CUG mistranslation stood as one of the highest among all samples. Additionally, we re-analysed our own dataset (wild-type T0 strain) using our pipeline with the “unbiased” reference database described in the manuscript (albeit without removing sequence redundancy). This involved extracting all proteins containing CUG sites from our diploid database and modifying them to incorporate each one of the 19 amino acids at CUG positions (excluding isoleucine mutations). By employing this approach, we found that 95.8% of CUG sites were indeed translated as serine, confirming the reassignment of serine for the CUG codon in *C. albicans,* and the remaining 4.2% of CUG sites were assigned to other codons, predominantly threonine, aspartic acid, glutamine, and leucine/isoleucine (Figure S8). However, it is important to exercise caution when interpreting these results. Firstly, certain amino acid substitutions may introduce a mass shift that could potentially be attributed to PTMs, such as methylated serine, offering an alternative explanation for Ser→Thr substitution. Secondly, the database used in this approach contains numerous entries with redundant regions, which increases the search space and may diminish statistical power. Lastly, employing an altered and incomplete proteome for the database search, where proteins lacking CUG sites were removed, can potentially lead to peptide misidentification, and introduce bias towards the expected mutations.

The findings presented in our study are thus in agreement with previous research conducted using various MS approaches, fluorescence, and *in vitro* tRNA charging methods (Suzuki *et al*., 1997, Gomes *et al*., 2007, Bezerra *et al*., 2013), which collectively refute the study by Mühlhausen *et al*. (Mühlhausen *et al*., 2021). Despite the low sensitivity and high variability among datasets and even between replicates from the same dataset, we found that whenever CUG mistranslation was identified, the associated error frequency was always higher than the global error of the sample (Figure 8). While we did observe some Ser-Leu substitutions associated with other serine codons (particularly UCA) in some samples, the error frequency was not meaningful. These results validated the specificity of Ser-Leu translation at CUG sites, as no other amino acid was detected in this codon.

**Figure 8.**
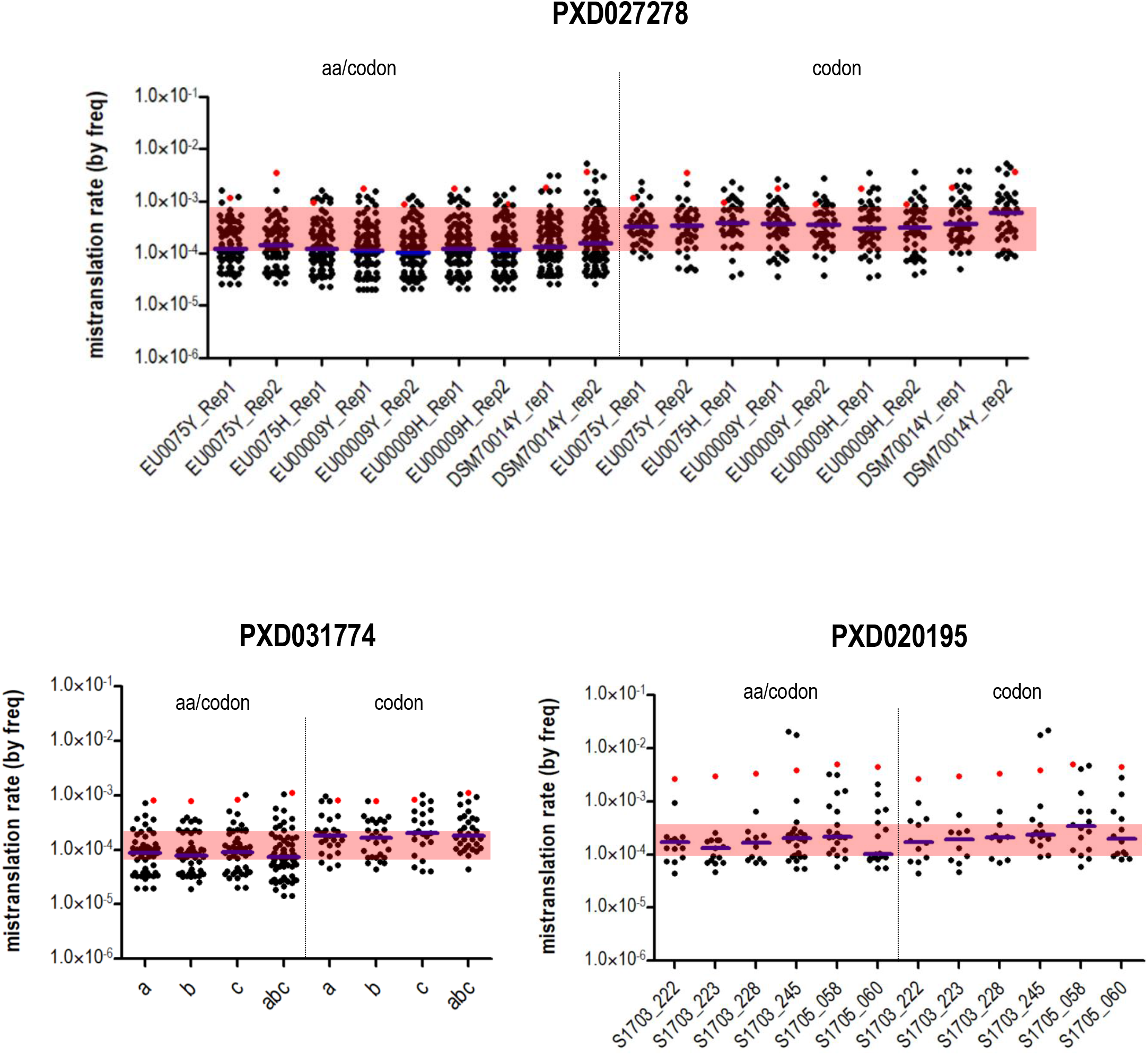
Global mistranslation analysis using datasets deposited on PRIDE repository. Distribution of mistranslation frequencies calculated for each codon mutation either specific for one single amino-acid (left) or for all detected substitutions (right). The blue line indicates the median of all calculated mistranslation frequencies. CUG codon mistranslation frequency is highlighted in red.

In the present study, we further demonstrate that protein biosynthesis errors in multiple codons are common in *C. albicans*, and that certain codons are more error-prone than others. Similar results were obtained in *Escherichia coli* and *Saccharomyces cerevisiae* by Mordret and co-workers who developed a pipeline for the identification of amino acid substitutions based on MaxQuant algorithms and the use of spectral libraries (Mordret *et al*., 2019). We reanalysed the *S. cerevisiae* sample of these studies (six fractions of the peptide mixture separated by strong cation exchange (Kulak *et al*., 2014)) and obtained a very similar result for the top 10 amino acid misincorporations (Figure S9). Although the type of amino acid misincorporations found in our study for *C. albicans* are substantially different, questioning the idea of a universal error pattern for mistranslation, there are many similarities with our own *S. cerevisiae* data set (Figure S10).

These carefully controlled studies demonstrate the utility of our pipeline in the study of protein biosynthesis errors although they also highlight the impact that different sample preparation procedures and thus different datasets have on the detection of amino acid substitutions and on the identity of substitution types, as well as on other proteomic outcomes (Mostovenko *et al*., 2013, den Ridder *et al*., 2022).

## Discussion

Over the past decade, the fields of proteomics and proteogenomics have seen significant improvements due to the development of more sensitive and high acquisition rate MS instruments, optimized software tools with user-friendly interfaces, and the availability of data in public repositories (Halder *et al*., 2021). While protein identification has been widely achieved through database search approaches requiring prior knowledge and availability of genome and/or proteome reference sequences (Verheggen *et al*., 2020), new strategies have been developed to detect novel peptides or peptide variants arising from protein isoforms absent from reference databases. To increase peptide identification, a common approach is the use of spectral libraries composed of previously observed and identified MS/MS spectra (Shao & Lam, 2017). Alternatively, *de novo* sequencing can be used to identify peptide-derived tandem mass spectra without prior knowledge of the sample or a pre-defined sequence database (Vitorino *et al*., 2020). While computationally intensive, *de novo* sequencing is unbiased and essential when no database is available. Search engines can also benefit from hybrid identification methods, combining *de novo* sequencing and database search to improve sensitivity and accuracy. This approach is exploited by PEAKS software which performs *de novo* sequencing of spectra before any database searching (Zhang *et al*., 2012). A complete and comprehensive reference database is critical, as incomplete databases may lead to unidentified or misidentified peptides resulting in false positive identifications (Knudsen & Chalkley, 2011). For unmatched *de novo* sequences, new algorithms can identify peptide variants to increase peptide identity (Han *et al*., 2005, Han *et al*., 2011).

Our goal was to detect and identify peptide mutations resulting from amino acid misincorporations in the human pathogenic fungus *C. albicans*, with the ambition of conducting a comprehensive analysis of protein biosynthesis errors in total protein extracts. To achieve this, we aimed to increase the sensitivity of our MS method while maintaining full codon representativeness. We opted to fractionate the samples using one-dimensional gel electrophoresis and obtain peptides via in-gel digestion of eight band segments for separate injection and independent sequencing analysis. This approach increased the dynamic range and depth of our analysis, allowing us to detect more peptide variants in complex mixtures (Shevchenko *et al*., 2006). To identify amino acid substitutions, we utilized PEAKS algorithms. We first performed a PEAKS PTM search before conducting a mutation search using SPIDER to avoid incorrectly matching of unexpected proteins or protein isoforms. We also included a list of common contaminants in our reference database and carefully selected protein isoforms resulting from allelic variations to prevent their misidentification as amino acid substitutions. However, the use of a reference database for proteogenomics has limitations, and some putative errors on codon assignment cannot be avoided. Although our strains are derived from the clinical isolate SC5314 whose sequence was used as a reference database, we cannot rule out the possibility that genetic variations may have occurred in our bioengineered strains that are not reflected at the proteome level. In addition, multiple peptide-protein matching to different proteins, different alleles or even within the same protein, with genetic different sequences, can also be an important source of error.

In this study, we show that by analysing the codon frequency of the peptides identified by MS/MS, we can obtain a relative quantification of codons-specific mistranslation at the proteome scale. However, the very low level of amino acid misincorporations detected remains a significant technical challenge to produce global maps of protein synthesis errors. In our analysis of *C. albicans* MS raw data available in the PRIDE archive, no Ser→Leu substitutions were found in most cases, even though up to three CUG-Leu incorporations were detected in some samples. This suggests that sample preparation, digestion and fractionation for protein synthesis error detection requires important optimization. For example, simple lysis buffer devoid of detergents (making extraction of membrane proteins inefficient), or a single digestion enzyme (producing extremely short or long peptides that are not detected or produce poor quality data) can create a systematic bias and generate false negatives in the acquired datasets (Kim *et al*., 2016). As Leu and Ser are chemically distinct amino acids, Ser→Leu substitutions may lead to rapid protein turnover both *in vivo* and during sample preparation, or to significant alterations in peptide behaviour inside the MS system, further complicating the detection of mutant peptides. This is consistent with our observation that 62% of the down-regulated proteins in T1 contain at least one CUG-encoded residue, potentially explaining why proteins in T1 are still identified, but their CUG-containing regions (peptides) are not.

The proteomic analysis conducted on a hypermistranslating strain (T1) supports the hypothesis that *C. albicans* tolerates very high levels of Leu incorporation at CUG sites, which is essential to explain the functional roles of Leu-incorporation on phenotypic diversity and pathogenesis (Bezerra *et al*., 2021). Our data also shows that Leu-misincorporation is specific to CUG codons and that its elevation affects the expression of proteins associated with ubiquitination and autophagy, and translation-related processes, highlighting its impact on protein processing and synthesis.

Interestingly, one of the Leu-CUG peptides belonged to Cdc60, which encodes the cytosolic leucyl tRNA synthetase (LeuRS). Notably, Cdc60 possesses a C-terminal domain where a CUG codon is localized, enabling it to recognize both the hybrid tRNA(CAG)Ser and its cognate tRNA(Leu) (Ji *et al*., 2016). Both the LeuRS-Ser and LeuRS-Leu isoforms catalyse activation and aminoacylation; however, the Leu isoform demonstrates higher activity (Zhou *et al*., 2013). Also, SerRS isoforms, another critical component involved in CUG decoding and containing its own CUG codon, exhibit similar characteristics (Rocha *et al*., 2011). To the best of our knowledge, this study provides the first evidence that both isoforms can coexist within live cells of *C. albicans*, thereby underscoring the importance of conducting a more comprehensive analysis to explore the full spectrum of CUG ambiguities.

In summary, our proteogenomics pipeline has successfully identified amino acid substitutions resulting from translational errors in the diploid fungus *C. albicans,* revealing varying levels of CUG mistranslation among different strains. This tool provides a valuable resource for assessing protein biosynthesis errors in various *C. albicans* strains, including clinical isolates, without the need of heterologous or synthetic reporters. The developed R scripts facilitate peptide filtration, codon assignment, and error frequency determination, while also preserving intermediate information on duplicates and putative errors in codon assignment for further in-depth analysis if required. For a comprehensive analysis of mistranslation events in *C. albicans,* we recommend whole-cell proteome fractionation by gel electrophoresis (>8 fractions) for separate trypsin digestion and MS/MS analysis, the use of comprehensive cell lysis buffer to ensure sample representativeness and long LC runs. Incorporating *de novo* sequencing to enhance peptide detection, utilizing the diploid *C. albicans* database to account for allelic variations, and considering PTMs mass shifts are all relevant due to the high noise-to-signal ratio of the analysis. The proposed pipeline and the multiple analyses conducted in this study with both our own MS/MS dataset and publicly available proteomics data, advance significantly our capacity to detect protein synthesis errors at the proteome level using mass spectrometry, however they also highlight the large challenges of obtaining complete maps of protein synthesis errors and, more importantly, how far we still are from having a methodology to comprehensively quantify amino acid misincorporations at this scale. Despite the technical difficulties, we anticipate that expected increases in MS sensitivity, development of new sample preparation methods and integration of artificial intelligence (AI) techniques in our pipelines will further enhance our capacity to explore in depth the translational misincorporation of amino acids into proteins (Chen *et al*., 2020). This is essential to better understand the biology of mistranslation.

## Materials and Methods

### Whole-cell lysis and protein extraction

*Candida albicans* cells from strains T0 (wild-type) and T1 (Bezerra *et al*., 2013) were grown at 30°C in YPD medium (2% glucose, 2% peptone, 1% yeast extract) for 16h or 22h, respectively, diluted at OD 0.1 in fresh medium and incubated at 30°C until OD 1.0-1.5. 20 mL of yeast suspension were harvested by centrifugation, washed 3x with PBS (137 mM NaCl, 2.7 mM KCl, 10 mM KH_2_PO_4_, 1.8 mM NaH_2_PO_4_, pH 7.4) and resuspended in lysis buffer (50 mM PBS pH 7.4, 1 mM EDTA, 5% glycerol, 1 mM PMSF and protease inhibitors from Roche). Cell lysis and protein extraction was achieved through mechanical lysis using glass beads and Precellys®24 Homogenizer (3 cycles of 30 seconds at 5500 rpm, alternating with 2 minutes on ice). Protein quantification was carried out using Pierce™ BCA Protein Assay Kit.

### Sample preparation and mass spectrometry

50 µg of extracted proteins were resolved on 12% SDS-PAGE. The gel was fixed (40% methanol, 10% acetic acid) for 30 minutes, stained with Coomassie blue for 1h at room temperature and destained with 25% methanol to remove the excess of dye. Protein bands were manually excised from the gel and transferred to Eppendorf tubes. Each sample/lane was divided into 8 fractions for separate enzymatic digestion and MS injection. Gel pieces were sequentially washed with 25 mM ammonium bicarbonate (AMBIC), then with 50% acetonitrile (ACN) in 25 mM AMBIC (as many times as needed to remove the dye), and once with ACN (30 min each wash). Cysteine residues were reduced with 10 mM DTT in 25 mM AMBIC (45 min at 56 °C) and alkylated with 55 mM iodoacetamide in 25 mM AMBIC (30 min at RT in the dark). A new round of washes (25 mM AMBIC, 50% ACN in 25 mM AMBIC for 15 min, and ACN for 10 min) was performed, and gel pieces were dried and later rehydrated in the digestion buffer. Proteins were digested as recommended by Shevchenko *et al*. (Shevchenko *et al*., 1996) with few modifications. Trypsin was added at an enzyme-to-substrate ratio of 1:50 (w/w) in 50 mM AMBIC. After 45 min on ice, the excess of buffer was removed, 50 mM AMBIC were added to cover the gel pieces, and the samples were incubated overnight at 37°C. Extraction of tryptic peptides was achieved by washing once with 5% formic acid (FA) and twice with 5% FA in 50% ACN (20 min each incubation). Samples were dried and resuspended in 1% FA. Each fraction of tryptic peptides was injected and separately analysed on an Ultimate 3000 HPLC system coupled to a QExactive Orbitrap (Thermo Fisher Scientific). The trap (5 mm × 300 µm) and the EASY-spray analytical (150 mm × 75 µm) columns used were C18 Pepmap100 (Dionex, LC Packings) with a particle size of 3 µm. Peptides were trapped at 30 μl/min in 96% solvent A (0.1% FA). Elution was achieved with the solvent B (0.1% FA / 80% ACN (v/v)) at 300 nl/min. The 92 min gradient used was as follows: 0–3 min, 96% solvent A; 3–70 min, 4–25% solvent B; 70–90 min, 25–40% solvent B; 90–92 min, 90% solvent B; 90–100 min, 90% solvent B; 101-120 min, 96% solvent A. The mass spectrometer was operated at 1.7 kV in the data-dependent acquisition (DDA) mode. A MS2 method was used with a FT survey scan from 400 to 1600 m/z (resolution 70000; AGC target 1E6). The 10 most intense peaks were subjected to HCD fragmentation (resolution 17500; AGC target 5E4, NCE 28%, max. injection time 100 ms, dynamic exclusion 35 s).

To analyse the impact of using exclusion lists between additional runs of MS/MS acquisition, raw data from the first MS/MS injection was analysed with Proteome Discoverer software and lists were generated (one for each fraction) comprising the most abundant peptides. The same sample was reinjected in the spectrometer using these exclusion lists (limited to 5000 peptides per run) to avoid analysis of redundant peptides.

### Mass spectrometry data analysis

Mass spectrometry raw data was processed by PEAKS Studio XPro software (Bioinformatics Solutions Inc., Waterloo, Ontario, Canada). For analysis, raw data from different fractions of the same sample were combined. Carbamidomethylation was set as a fixed modification, and oxidation of methionine (M), deamidation (NQ) and protein N-acetylation were set as variable modifications. Searches were performed using mass tolerances of 5 ppm for precursor ions and 0.02 Da for fragment ions. Trypsin digestion was set as specific with up to 2 missed cleavages allowed. Sampleś raw data was initially searched against a haploid reference proteome database, obtained from the *C. albicans* diploid genome assembly 22 (Muzzey *et al*., 2013, Skrzypek *et al*., 2017) and available at Candida Genome Database (CGD), where a single allele represents each pair in the diploid genome: *C_albicans_SC5314_version_A22-s07-m01-r149_default_protein.fasta*, plus a list of common proteomics contaminants obtained from the common Repository of Adventitious Proteins (cRAP) (Mellacheruvu *et al*., 2013). The significance of protein/peptide identifications was controlled to 1% FDR (*False Discovery Rate*) at both peptide and protein level. The list of peptides and identified proteins (*protein-peptides* and *proteins* files – Supplementary data) was used to evaluate proteome and protein coverage and analyse the number and frequency of each codon type assigned to the identified proteins and peptides in the samples. To assess *C. albicans* global mistranslation, the same raw data was searched against the diploid reference proteome database available also at CGD which contains sequence information from both haplotypes A and B: *C_albicans_SC5314_version_A22-s07-m01-r149_orf_trans_all.fasta*, plus the cRAP list. To reduce database size and redundancy, we removed the haplotype B from protein duplicates with identical amino acid sequences, and 20 entries from proteins encoded by mitochondrial genes (Figure S3). Unspecified post-translation modifications (PTMs) were uncovered by PEAKS PTM algorithm in which all PTMs were selected except those marked as “Isotopic label” or “Chemical derivatives”. The remaining unmatched spectra were analysed by SPIDER algorithm to identify amino acid substitutions. The significance of protein/peptide identifications was controlled to 1% FDR at both peptide and protein levels. PTMs localization accuracy was supported by AScore, meaning that the probability that the modification occurred at the reported position compared to other possible positions was at least 0.01. To confirm amino acid substitutions, a pair of b or y ions was found with at least 5% relative intensity showing fragmentation before and after the modified amino acid. Thus, peptides with at least one PTM above the Ascore (depicted in the *PTM* column at *protein-peptides* file – Supplementary data) and peptides with at least one substitution above 5% Ion Intensity (selected from the *peptides* list containing only mutations found by SPIDER – Supplementary data) were validated and maintained in the dataset. Peptides with insertions and deletions were also validated by the ion intensity filter. Finally, peptides with robust amino-acid substitutions were further filtered according to the existence of their unmodified counterpart, using R scripts. Redundant peptide sequences due to allelic duplicates or paralogs (with same m/z and retention time) were also removed from the final dataset.

### Error frequency determination

*C. albicans* genome sequence was obtained from CGD: *C_albicans_SC5314_version_A22-s07-m01-r149_orf_coding.fasta* which matched the diploid proteome database used for reference. Codons were assigned to amino acids from all validated peptides identified by the different algorithms from PEAKS software, using R scripts. The frequency of all codons was assessed, and the mistranslation error was calculated for each specific amino acid substitution / codon pair and for each codon independently on the destination amino acid, using the frequency of mutated codons per the frequency of all identified codons present in the final dataset. For example, for CUG codon: [frequency of CUG codons translated to other amino acids besides serine / frequency of all detected CUG codons in the sample, independently on the incorporated amino acid] x 100. For error frequency assessment we disregarded substitutions indistinguishable from PTMs or artefacts in the Unimod database {http://www.unimod.org}: Ala→Ile/Leu (tri-Methylation); Asp→Glu (Methylation); Phe→Tyr (Oxidation); Asn→Asp/Gln (Deamidation/Methylation); Pro→Glu (di-Oxidation); Gln→Glu (Deamidation); Ser→Asp/Glu/Thr (Formylation/Acetylation/Methylation); Thr→Glu (Formylation).

### Proteome coverage and codon frequency analysis

Proteome and protein sequence coverage and peptide/protein abundance information were obtained from *proteins* file (Supplementary data) exported after PEAKS DB search against the “haploid” *C. albicans* database. The area under the curve of the peptide feature found at the same m/z and retention time as the MS/MS scan (specified in Area column) was used as an indicator of peptide and protein abundance. The list of identified peptides (*protein-peptides* file – Supplementary data) was filtered according to the parameters described before and assigned to their codons using the matching genome sequence available at CGD: *C_albicans_SC5314_version_A22-s07-m01-r149_default_coding.* Global codon frequency and relative synonymous codon usage (RSCU) were assessed for all validated peptides, after PEAKS DB search against the “haploid” *C. albicans* database, using R scripts. For this, peptide variants (such as identical peptide sequences but with different PTMs or in different locations) and peptide duplicates due to multiple protein matching, were excluded from the dataset. The RSCU from all identified proteins was obtained by Anaconda software using the non-standard translation table for alternative yeast codon usage (Pinheiro *et al*., 2006). Protein CUG content was obtained from CGD and gene ontology analysis was performed using the GO Term Finder at the CGD website. Venn diagrams were obtained with the FunRich 3.1.3 program.

### Label-free quantification

An ID-directed label-free quantification (LFQ) provided by PEAKS Q module in PEAKSXpro software was used to compare the abundance of proteins from T0 and T1 strains. This approach is based on the relative abundance (MS1 feature area) of all identified peptide features detected in the two samples. Data were normalized to the total ion current (TIC) and filtered according to quality (≥10), significance (≥20) and fold change (≥2). A volcano plot was obtained by PEAKS software and information regarding protein area (top-3 peptides) and the ratio between T0 and T1 was extracted from *proteins.csv* file.

### Flow cytometry

T0 and T1 strains expressing different variants of GFP (Bezerra *et al*., 2013) were grown at 30°C in YPD medium for 16h or 22h, respectively, diluted at OD 0.15 in fresh medium and incubated at 30°C until OD 1.0-1.5. Cells were recovered in phosphate-buffered saline (PBS) at 1:10 dilution, washed once and filtered (0.45 µm) before being analysed in a BD Accuri™ C6 Flow Cytometer. Fluorescence profiles (FL1) of 20000 events for each sample were collected only on those cells appearing in R1 to exclude agglutinates and debris. The fluorescence intensity of each GFP variant was analysed: GFP-Leu_201_ (positive control); GFP-Ser_201_ (negative control); GFP-Ser/Leu_201_ (CUG reporter). The assay was replicated three times and Leucine incorporation was calculated for each one according to the formula: [(GFP-Ser/Leu_201_) – (GFP-Ser_201_)] / [(GFP-Leu_201_) – (GFP-Ser_201_)].

## Data and code availability

Raw data were analysed with PEAKS XPro software, and the resulting files were processed using a custom pipeline written in R. All data will be made available upon request.

## Supporting information

peptide

protein-peptides

proteins

