## Supplemental figures for "A proteogenomic pipeline for the analysis of protein biosynthesis errors in the human pathogen *Candida albicans*"

### Supplementary data

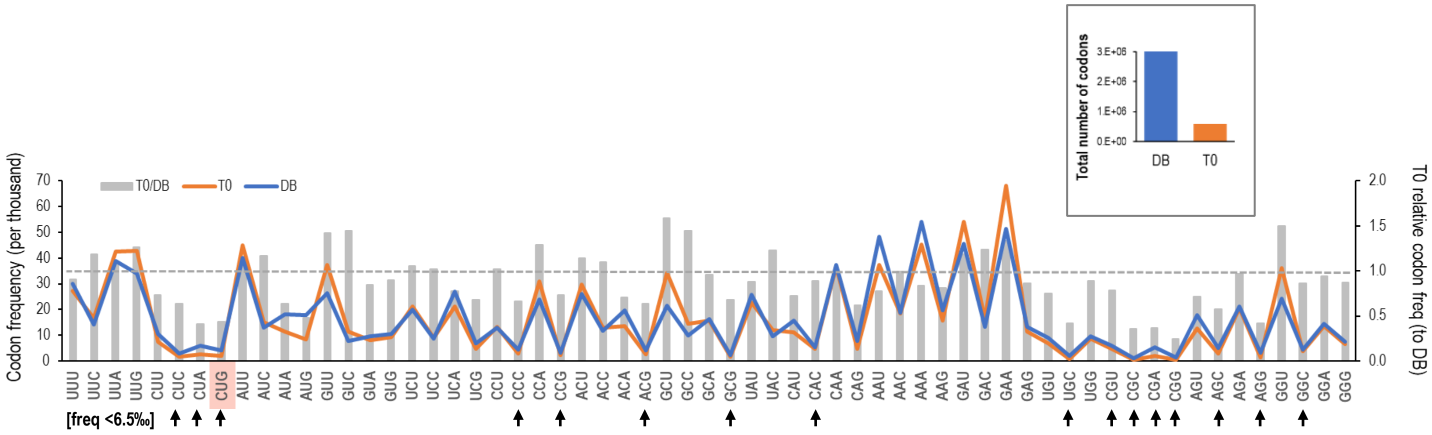


**Figure S1.** Codon frequency analysis. Lines in the graphic compare the *C. albicans* codon frequency obtained for the reference proteome used for PEAKS DB search and calculated by Anaconda software, with the calculated codon frequency for the T0 sample after codon assignment to all identified peptides (without duplicates). The columns highlight codon frequency deviations regarding the reference DB, showing the ratio of T0 codon frequency to the one obtained for the reference. The dashed line indicates the expected ratio if no differences were observed. Arrows point to rare codons, defined as those whose frequency is lower than half of the median of all codon frequencies obtained for the DB used for reference (6.5 codons per thousand). The box shows the total number of codons used for this analysis.

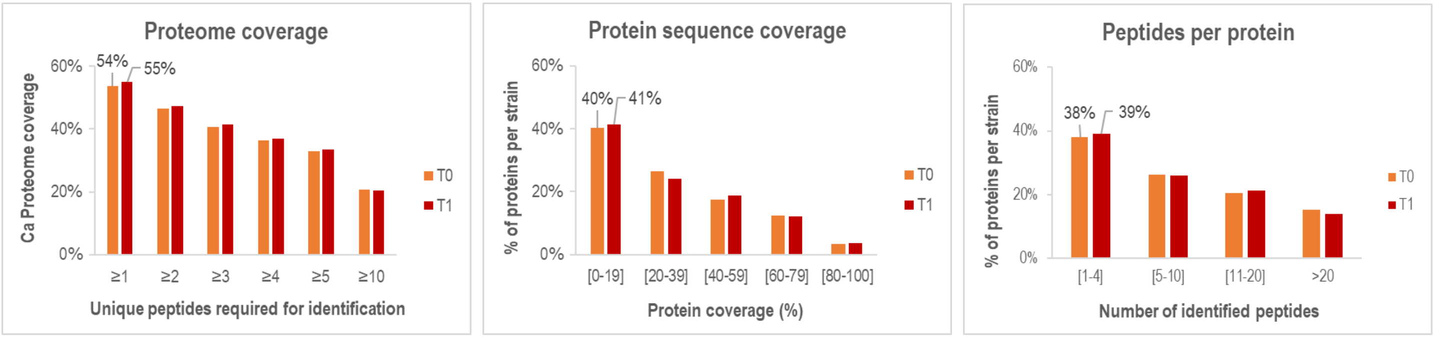


**Figure S2**: Percentage of proteome coverage, sequence coverage and identified peptides per protein, for T0 and T1 samples.

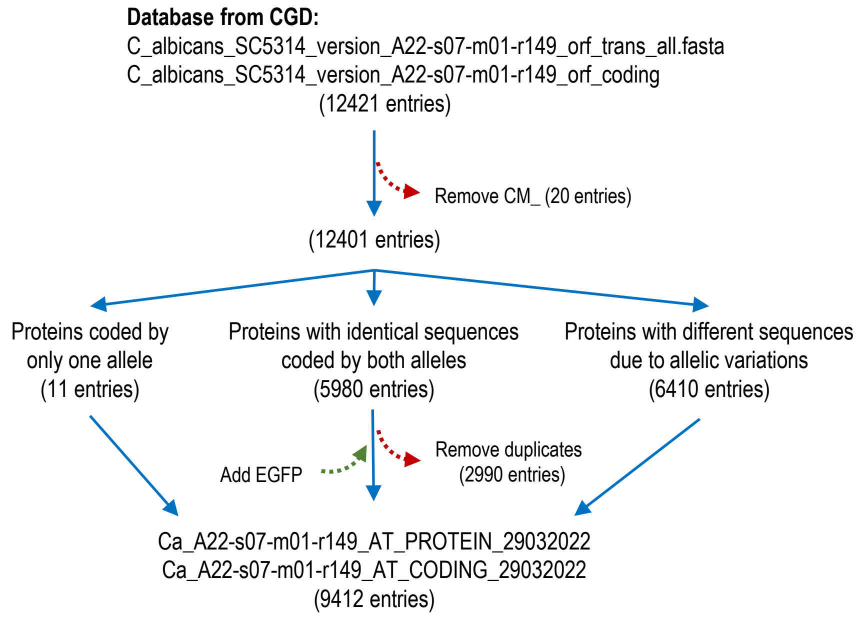


**Figure S3**: *C. albicans* diploid proteome database used for Database Search algorithm.

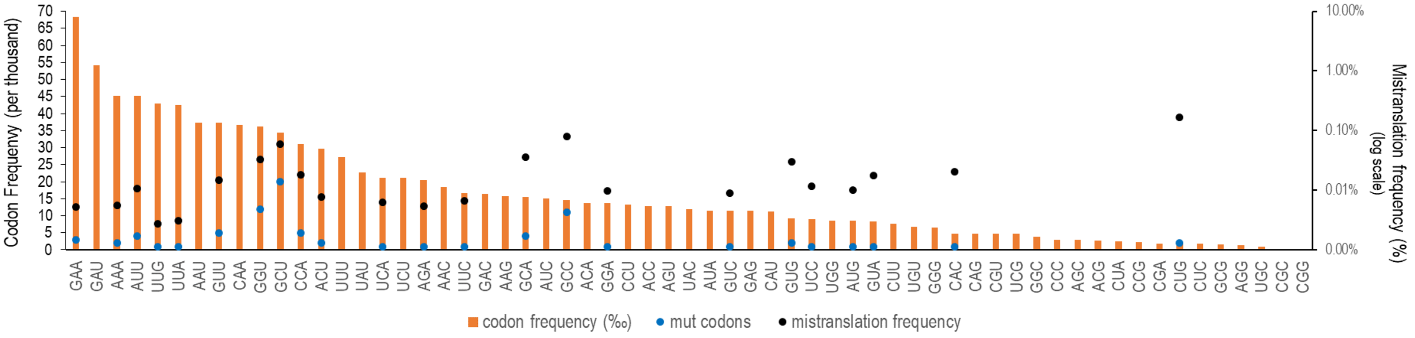


**Figure S4**: Relation between codon frequency and mistranslation frequency.

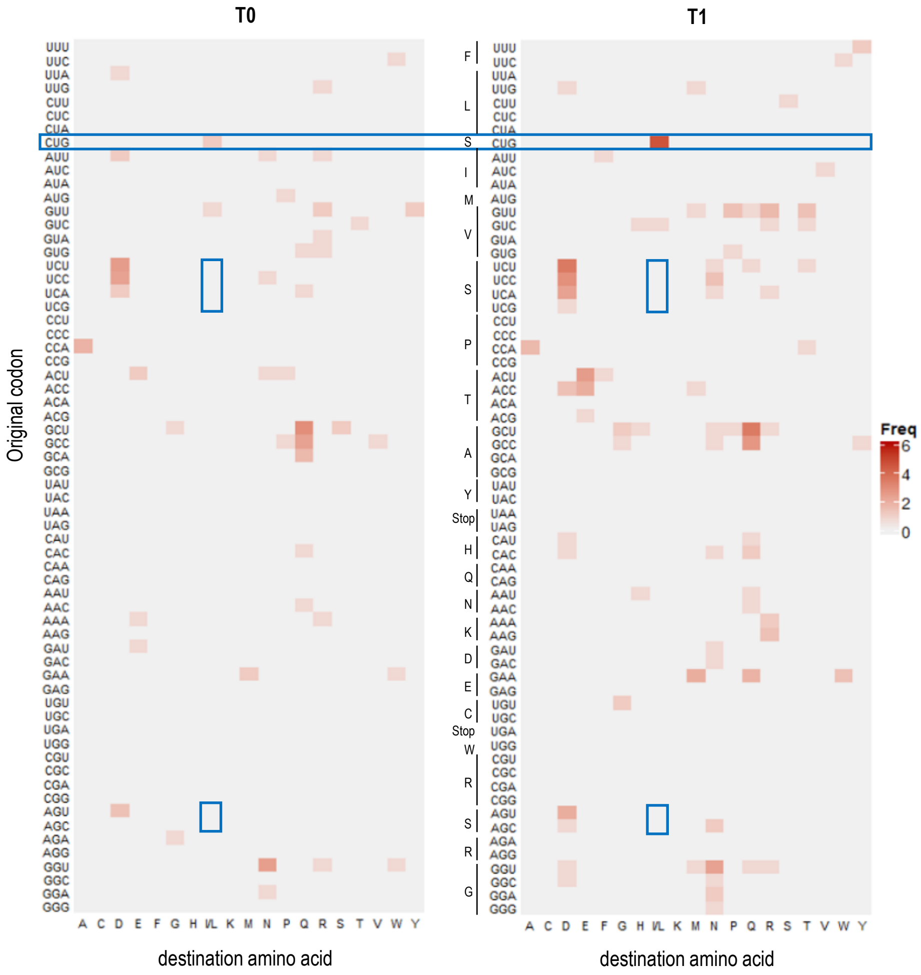


**Figure S5.** Matrix of the number and identity of amino acid substitutions in T0 and T1 strains. Highlighted in blue boxes are amino acid substitutions in the CUG codon and Ser→Leu substitutions in serine codons.

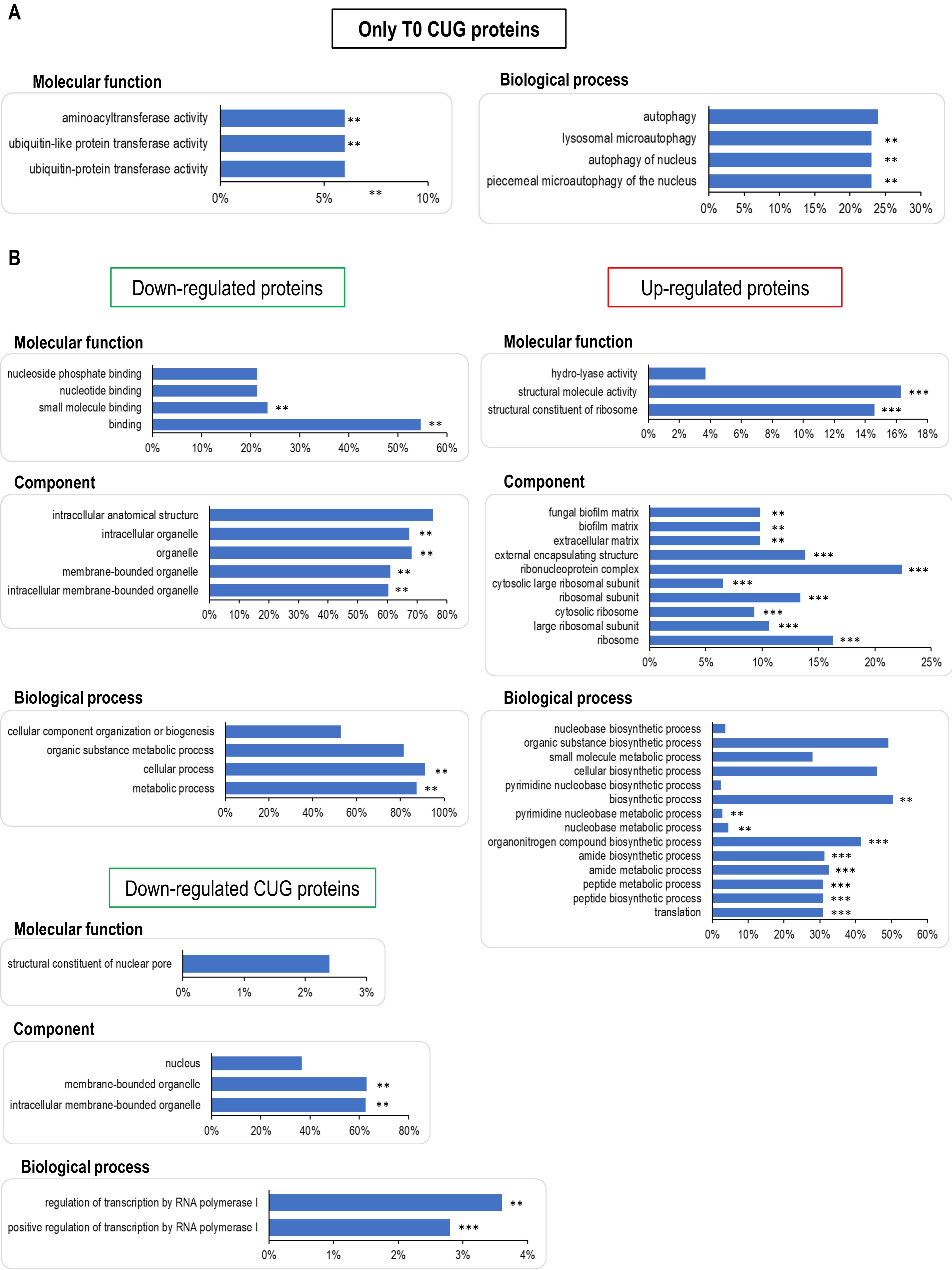


**Figure S6. A)** GO enrichment analysis of CUG-containing proteins exclusively found in the T0 sample (background: 3680 proteins identified in both strains). **B)** GO enrichment analysis of T1 up and down regulated proteins (background: 3063 common proteins shared by both strains). GOTermFinder application at CGD. p-value cut-off: 0.05. **p<0.01; ***p<0.001.

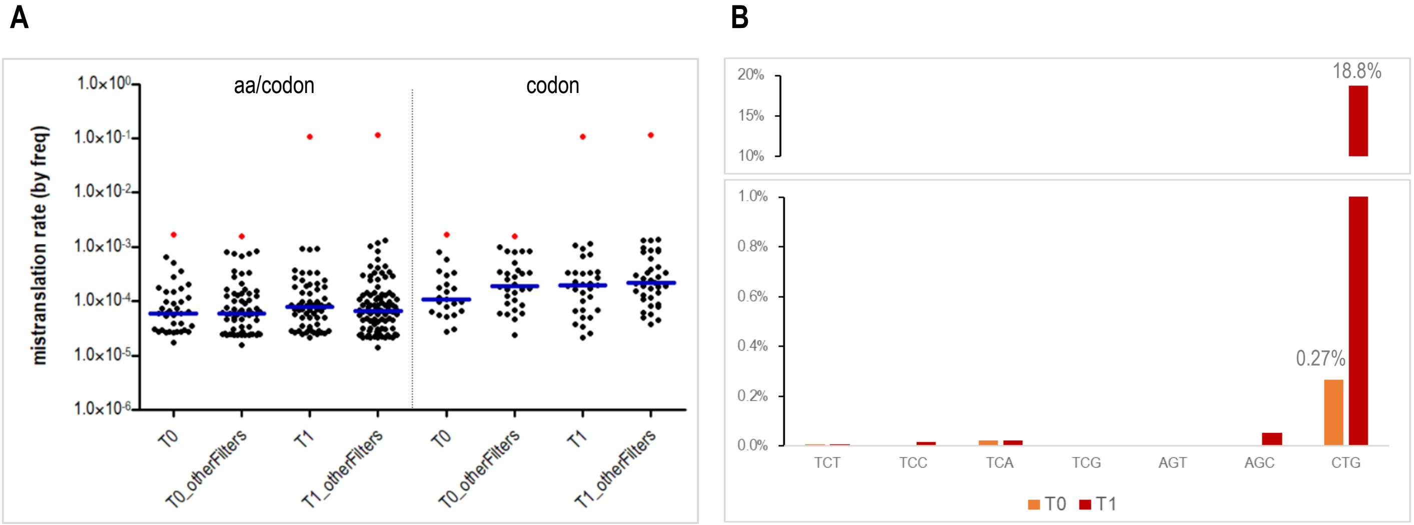


**Figure S7.** Impact of using more lenient filters (_otherFilters) **(A)** and Ser→Leu directed searches **(B)** in error frequency determination. Data was acquired and analysed as described in Material and Methods with a few modifications: **A)** searches were performed using mass tolerances of 10 ppm for precursor ions, trypsin digestion was set as semi-specific and 1% ion intensity was used to validate amino acid substitutions; **B)** Ser→Leu substitution [S(+26.05)] was additionally set as variable PTM for PEAKS DB search, and the list of peptides was obtained from the *protein-peptides.cvs* file exported after DB search, without running PEAKS PTM and SPIDER algorithms. **A)** Distribution of mistranslation frequencies calculated for each codon mutation either specific for one single amino-acid (left) or for all detected substitutions (right). The blue line indicates the median of all calculated mistranslation frequencies. L(S)CUG mistranslation frequency is highlighted in red. **B)** Comparison between the mistranslation frequencies calculated for serine codons due to Ser→Leu substitutions from T0 and T1 strains when performing a directed search.

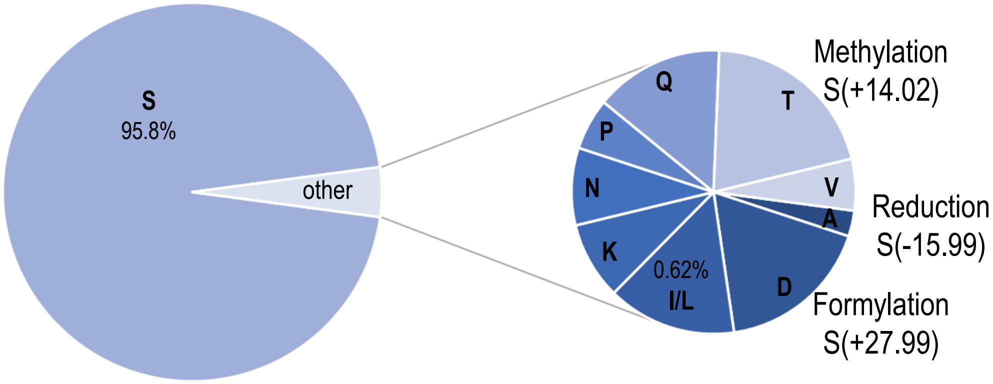


**Figure S8.** Pie chart showing the identity of amino acids assigned to the CUG codon in a wild-type strain (T0), when using a mutated database as reference for PEAKS DB search. This mutated database was generated using R scripts and composed only by CUG containing proteins.

**A**

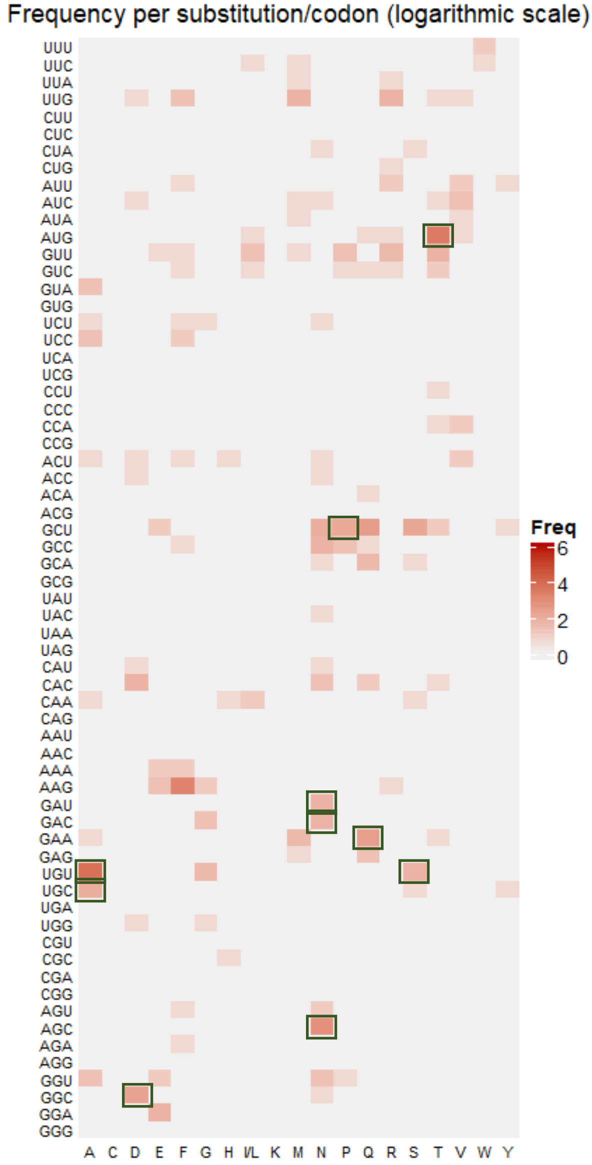


**B**

| Most frequent substitutions using our pipeline | | Frequency |
|---|---|---|
| A(C) | TGT | 41 |
| T(M) | ATG | 31 |
| F(K) | AAG | 23 |
| N(S) | AGC | 16 |
| Q(A) | GCT | 11 |
| Q(E) | GAA | 10 |
| D(G) | GGC | 9 |
| S(A) | GCT | 8 |
| P(A) | GCT | 7 |
| A(C) | TGC | 6 |
| N(A) | GCT | 6 |
| N(D) | GAC | 5 |
| N(D) | GAT | 5 |
| N(A) | GCC | 5 |
| E(G) | GGA | 5 |
| S(C) | TGT | 5 |
| T(V) | GTT | 5 |
| D(H) | CAC | 5 |
| R(L) | TTG | 5 |
| M(L) | TTG | 5 |

**C**

| Most frequent substitutions in original data (Mordret, et al., 2019) | |
|---|---|
| T(M) | ATG |
| A(C) | TGT |
| D(G) | GGC |
| N(S) | AGC |
| S(C) | TGT |
| A(C) | TGC |
| N(D) | GAT |
| N(D) | GAC |
| Q(E) | GAA |
| P(A) | GCT |

**Figure S9. A)** Matrix of the number and identity of amino acid substitutions in *S. cerevisiae* (SCX data) from PXD000269 project obtained from PRIDE repository. Amino acid substitutions indistinguishable from PTMs or artifacts in the Unimod database were removed from the dataset, as in Mordret, *et al.*, 2019. **B)** Most frequent substitutions found by our pipeline in frequency order (most frequent substitutions at the top). New substitution types are highlighted in grey. **C)** Top 10 substitutions found originally by Mordret and co-workers ordered by frequency (highlighted in green boxes at the matrix).

**Note:** with our pipeline we uncovered 86 new substitution types but missed 37 previously described by Mordret and co-workers. Nevertheless, the most frequent substitutions were mainly the same for both analysis.

**A**

| Most frequent substitutions using PRIDE Sc data and our pipeline | |
|---|---|
| A(C) | TGT |
| T(M) | ATG |
| F(K) | AAG |
| N(S) | AGC |
| Q(A) | GCT |
| Q(E) | GAA |
| D(G) | GGC |
| S(A) | GCT |
| P(A) | GCT |
| A(C) | TGC |

| Most frequent substitutions using our Sc data and pipeline | |
|---|---|
| Q(A) | GCT |
| Q(A) | GCC |
| N(G) | GGT |
| R(V) | GTT |
| E(K) | AAA |
| Q(E) | GAA |
| Q(A) | GCA |
| A(V) | GTT |
| T(V) | GTT |
| M(E) | GAA |

| Most frequent substitutions using our Ca data and pipeline | |
|---|---|
| Q(A) | GCT |
| N(G) | GGT |
| Q(A) | GCC |
| A(P) | CCA |
| Q(A) | GCA |
| D(I) | ATT |
| L(S) | CTG |
| M(E) | GAA |
| S(A) | GCT |
| R(V) | GTT |

**B**

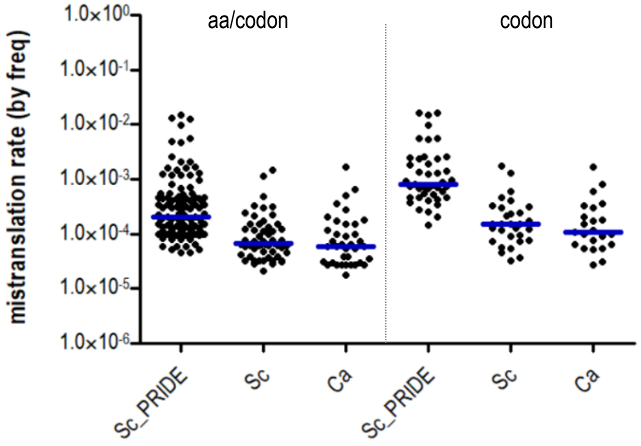


**Figure S10.** Comparative analysis of protein biosynthesis errors in different datasets: *S. cerevisiae* data obtained by SCX (PXD000269); our own *S. cerevisiae* data from 4 fractions separated by SDS-PAGE; and our *C. albicans* data from 8 fractions. **A)** Most frequent substitutions found by our pipeline in frequency order (most frequent substitutions at the top). **B)** Distribution of mistranslation frequencies calculated for each codon mutation either specific for one single amino-acid (left) or for all detected substitutions (right). The blue line indicates the median of all calculated error frequencies.
